# Protein ubiquitylation is essential for the schizont to merozoite transition in *Plasmodium falciparum* blood-stage development

**DOI:** 10.1101/755439

**Authors:** Yang Wu, Vesela Encheva, Judith L. Green, Edwin Lasonder, Adchara Prommaban, Simone Kunzelmann, Evangelos Christodoulou, Munira Grainger, Ngoc Truongvan, Sebastian Bothe, Vikram Sharma, Wei Song, Irene Pinzuti, Chairat Uthaipibull, Somdet Srichairatanakool, Veronique Barault, Gordon Langsley, Hermann Schindelin, Benjamin Stieglitz, Ambrosius P. Snijders, Anthony A. Holder

## Abstract

Ubiquitylation is a common post translational modification of eukaryotic proteins and in the human malaria parasite, *Plasmodium falciparum (Pf)* overall ubiquitylation increases in the transition from intracellular schizont to extracellular merozoite stages in the asexual blood stage cycle. Here, we identify specific ubiquitylation sites of protein substrates in three intracellular parasite stages and extracellular merozoites; a total of 1464 sites in 546 proteins were identified (data available via ProteomeXchange with identifier PXD014998). 469 ubiquitylated proteins were identified in merozoites compared with only 160 in the preceding intracellular schizont stage, indicating a large increase in protein ubiquitylation associated with merozoite maturation. Following merozoite invasion of erythrocytes, few ubiquitylated proteins were detected in the first intracellular ring stage but as parasites matured through trophozoite to schizont stages the extent of ubiquitylation increased. We identified commonly used ubiquitylation motifs and groups of ubiquitylated proteins in specific areas of cellular function, for example merozoite pellicle proteins involved in erythrocyte invasion, exported proteins, and histones. To investigate the importance of ubiquitylation we screened ubiquitin pathway inhibitors in a parasite growth assay and identified the ubiquitin activating enzyme (UBA1 or E1) inhibitor MLN7243 (TAK-243) to be particularly effective. This small molecule was shown to be a potent inhibitor of recombinant PfUBA1, and a structural homology model of MLN7243 bound to the parasite enzyme highlights avenues for the development of *P. falciparum* specific inhibitors. We created a genetically modified parasite with a rapamycin-inducible functional deletion of *uba1;* addition of either MLN7243 or rapamycin to the recombinant parasite line resulted in the same phenotype, with parasite development blocked at the late schizont stage. These results indicate that the intracellular target of MLN7243 is UBA1, and this activity is essential for the final differentiation of schizonts to merozoites. The ubiquitylation of many merozoite proteins and their disappearance in ring stages are consistent with the idea that ubiquitylation leads to their destruction via the proteasome once their function is complete following invasion, which would allow amino acid recycling in the period prior to the parasite’s elaboration of a new food vacuole.

## Introduction

The malaria parasite’s life cycle includes an asexual blood stage infection that is responsible for the disease. In the case of the major human parasite, *Plasmodium falciparum*, this stage lasts approximately 48 hours and includes a very short-lived extracellular merozoite and an intra-erythrocytic development and multiplication phase. Following erythrocyte invasion by the merozoite, the parasite develops through haploid ring and trophozoite stages and then, in the schizont stage, DNA synthesis, mitosis and nuclear division lead to a multinucleate syncytium. At the end of schizogony, cytokinesis leads to the differentiation of new merozoites, which are subsequently released by rupture of the host erythrocyte (egress) and go on to invade other erythrocytes. During schizogony cellular structures are elaborated, including secretory organelles such as the micronemes, rhoptries and dense granules, and the inner membrane complex (IMC) that forms part of the merozoite surface pellicle. The parasite also profoundly modifies its host cell; it resides within a parasitophorous vacuole, but secretes numerous proteins into the cytoplasm, and extensively modifies the cytoskeleton and surface, of the erythrocyte. For each cycle of development, structures of the intracellular parasite such as the food vacuole are discarded at egress, and the merozoite secretory organelles are disassembled following invasion. Although it is difficult to distinguish morphologically extracellular merozoites from intracellular merozoites at the final stage of schizogony, there are substantial differences in the merozoite and schizont phosphoproteomes [1, 2], and a significant increase in protein ubiquitylation in extracellular merozoites [2].

Protein ubiquitylation plays key roles in cell biology, for example in tagging proteins for proteasomal degradation, or targeting to other distinct subcellular locations. Ubiquitin is a 76-residue protein that is usually added through a covalent isopeptide bond to the ε-amino group of lysine residues in substrate proteins. In a three step process the C-terminal carboxyl group of ubiquitin is activated by E1 (ubiquitin activating enzyme, UBA1) and then ubiquitin is transferred as a thioester to an E2 (ubiquitin conjugating enzyme) prior to transfer to substrate, with the assistance of an E3 (ubiquitin ligase). Ubiquitin can be removed from substrates by deubiquitinating enzymes (DUBs). Although single ubiquitin moieties may be added to a substrate protein, polyubiquitylation is also very common, in which the ubiquitylation of bound ubiquitin results in long ubiquitin chains.

Recently, for example in mammalian and plant cells, the ubiquitome has been examined using a method in which proteins are first digested with trypsin, to produce peptides containing a di-Gly remnant derived from the ubiquitin C-terminus (^74^Arg-Gly-Gly^76^) attached to the ubiquitylated residue [3, 4]. These peptides are purified using an antibody specific for the remnant, fractionated by liquid chromatography and then identified by mass spectrometry [4]. In this way, ubiquitylation sites and the corresponding substrates can be identified. This approach has been used to identify ubiquitylated proteins in the tachyzoite stages of *Toxoplasma gondii* [5], an apicomplexan parasite that is a distant relative of *P. falciparum*.

In the malaria parasite very little is known about the extent and role of protein ubiquitylation. Bioinformatic analyses have revealed genes encoding the necessary machinery in the parasite genome [6–8]. This machinery is located in two distinct subcellular compartments within the parasite: the cytoplasm and the apicoplast [9, 10]. Ubiquitylation substrates have been identified either by bioinformatic prediction, by mass spectrometry-based analysis of proteins purified by ubiquitin-affinity [11], or using tandem ubiquitin binding entities (TUBEs)-based methodologies [12, 13]. In one study 72 putative substrates were identified [11], and in a second study 12 proteins binding to TUBEs were detected [12]. However, the direct identification of ubiquitylated lysine residues in peptides diagnostic for substrate proteins has been used only in a very limited way while extracellular merozoites were not examined in any of the earlier studies.

In humans, four genes encode ubiquitin (Ub): two that code for precursors containing tandem repeats of ubiquitin and two that code for a single ubiquitin unit with C-terminal extensions encoding the 60S ribosomal protein L40 and the 40S ribosomal protein S27a, respectively. In the *P. falciparum* genome there is a single gene (PF3D7_1211800) encoding a polyubiquitin precursor with five copies of ubiquitin [14], and a putative L40 orthologue (PF3D7_1365900); in the putative S27a orthologue (PF3D7_1402500), the encoded N-terminal sequence differs substantially from that of ubiquitin. The sequences of *P. falciparum* and human ubiquitin are identical except for an Asp/Glu substitution at residue 16. The ubiquitin sequence has seven lysine residues at positions 6, 11, 27, 29, 33, 48 and 63, and each of these, as well as the N-terminal α-amino group, is a potential ubiquitylation site. When ubiquitin of either host or parasite origin is digested with trypsin, diGly-remnant peptides cannot be distinguished, except for those modified at position 11 or 27 for which the Asp/Glu polymorphism differentiates host- and parasite-derived sequences. Therefore, it is difficult to distinguish parasite- or host-derived polyubiquitylation products for an organism living in human erythrocytes.

Whilst there has been considerable interest in targeting the proteasome with newly identified drugs [15], until recently [16, 17] ubiquitin biology has been largely neglected in malaria research. There is extensive current research to identify inhibitors of the ubiquitin pathway for other therapeutic applications, particularly in the area of cancer chemotherapy [18]. One class of E1 inhibitor is the adenosyl sulphamates that form a covalent adduct with ubiquitin. This reaction is catalysed by E1 and inhibits further ATP-based activation of ubiquitin. MLN7243 (TAK-243) is one of these compounds; it inhibits the human (*Homo sapiens*) and yeast (*Saccharomyces cerevisiae*) ubiquitin E1 (HsUBA1 and ScUBA1) and structural data have been obtained for E1 with bound inhibitor and bound ubiquitin [18–20]. However, it is unknown whether MLN7243 inhibits *P. falciparum* (Pf) UBA1. Few other inhibitors of the ubiquitin pathway have been tested as malaria parasite inhibitors [21].

To validate the on-target specificity of antimalarial compounds it is now possible to use genetic approaches. Application of CRISPR-Cas9 technology to modify the parasite genome, allows genes to be deleted using an inducible Cre-recombinase system [22–24], or single codon changes conferring drug resistance to be introduced [25].

In this study we compared the ubiquitylated proteins of extracellular merozoites and intracellular stages, and confirmed that ubiquitylation is much more extensive in merozoites [2]. We show that some commercially available inhibitors of ubiquitylation are able to inhibit parasite growth, with the most active being MLN7243, which we show binds to the active site of PfUBA1 and inhibits its activity. We engineered a parasite line in which truncation of the UBA1 gene could be induced to produce a functional knockout. This modified parasite stopped development at the same point as treatment with MLN7243: the schizont to merozoite transition. The data indicate that ubiquitylation plays an essential role in parasite development and in the biology of the short-lived extracellular stage of the malaria parasite’s life cycle. These results greatly extend our knowledge of protein ubiquitylation in the malaria parasite and validate UBA1 as a target for further drug development.

## Results

### The *P. falciparum* ubiquitome: distribution of ubiquitylation sites and stage specificity

To identify ubiquitylation sites and substrate proteins in both intracellular and extracellular blood stage parasites, di-Gly remnant peptides produced by trypsin digestion of cell lysates were immunopurified and analysed by liquid chromatography-tandem mass spectrometry. Ubiquitylated peptides from both parasite and host proteins were identified by searching Q Exactive MSMS spectra against a combined *P. falciparum* and *H. sapiens* protein database using MaxQuant [26]. In total, we identified 1,464 distinct ubiquitylation sites in 546 parasite proteins (1% false discovery rate [FDR], site localisation probability >0.75), and 281 ubiquitylation sites in 115 host substrates (Figure 1, Supplementary Tables 1 and 2). 265 parasite proteins (48%) were represented in the ubiquitome by a single diGly peptide and approximately 90% (494) of the protein substrates contained between one and five ubiquitylation sites. The remaining proteins had between six and 25 experimentally detected ubiquitylation sites (Supplementary Table 3). These data extend considerably the number of identified ubiquitylation sites and substrate proteins from the small number (seventeen sites in five substrates) described previously [12]. Analysis of the sequences around the ubiquitylation sites identified 11 abundant motifs (representing 644 sites), each comprised of 13 amino acids with the modified lysine at the centre (Supplementary Figure 1).

**Figure 1.**
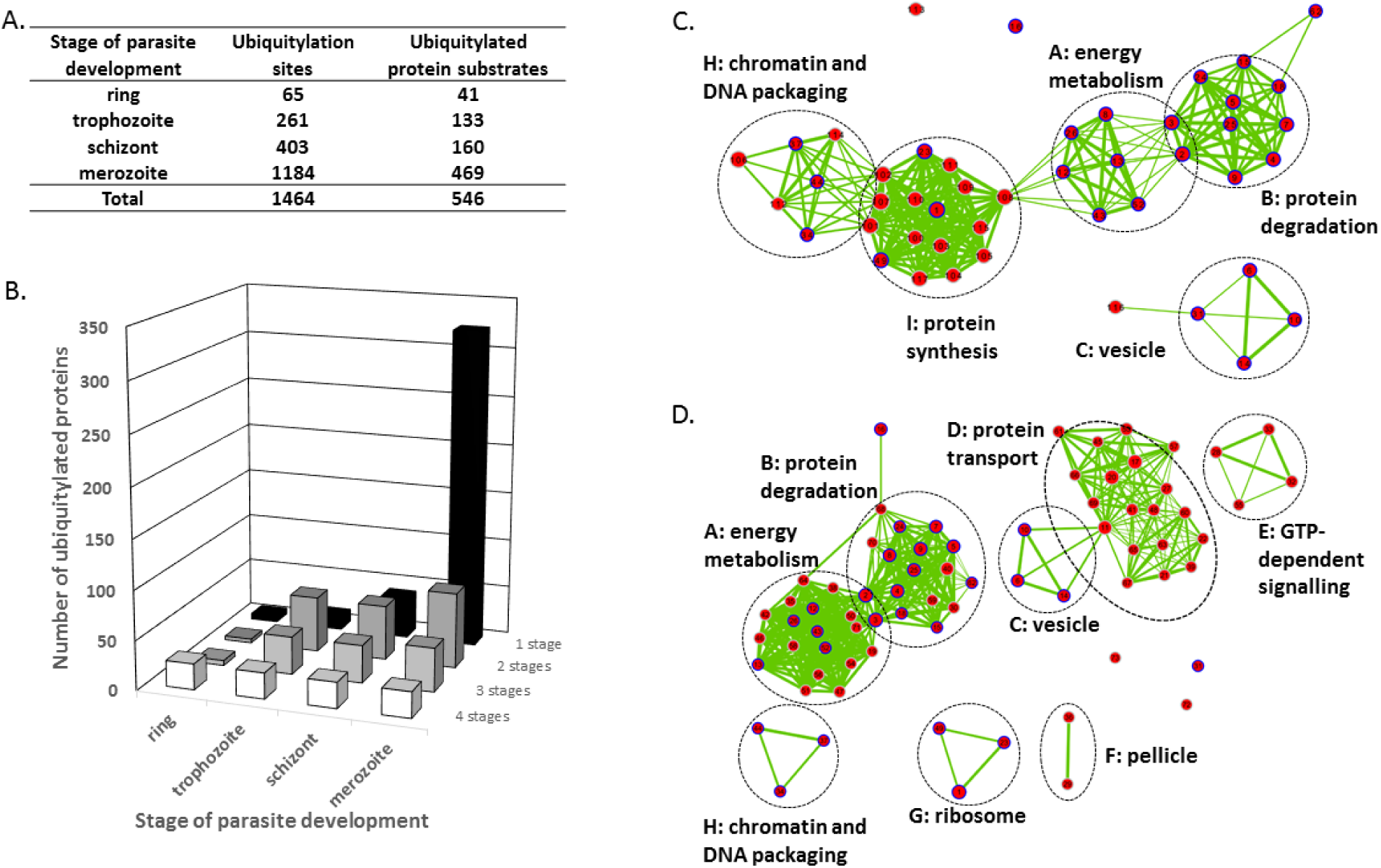
Ubiquitylation throughout the parasite asexual blood stage. (A) Ubiquitylation sites and their corresponding proteins in four stages of parasite development. (B) Distribution of ubiquitylated proteins between the intra-erythrocytic ring, trophozoite and schizont stages and the extracellular merozoite stage. Whilst some proteins are ubiquitylated in all four stages, the majority were found to be ubiquitylated only in merozoites. The primary data are presented in Supplementary Tables S1 and S3. Maps depicting GO enrichment results for (C) the intra-erythrocytic parasite and (D) the merozoite ubiquitomes. The network representation of enriched GO terms reveals large clusters (A-I) of functionally-related GO terms. Nodes with border lines in blue represent GO terms that are found enriched in the ubiquitome of all stages. Identification of the GO terms and associated proteins is provided in Supplementary Table 4.

Protein ubiquitylation (both the number of ubiquitylation sites and the number of protein substrates) increased during development of the intracellular parasite from rings through trophozoites to schizonts, but the largest increase in ubiquitylation was in extracellular merozoites (Figure 1 and Supplementary Tables 1 and 2). This substantial increase in the transition from schizonts to merozoites was followed by a substantial decrease in the transition to rings, the first intracellular stage. For some substrates certain sites were ubiquitylated in one stage of the cycle but not in others. Only 65 sites, representing 41 protein substrates, were detected in ring stages, and this increased to 261 and 403 sites representing 133 and 160 substrates in trophozoites and schizonts, respectively. However, 1184 di-Gly remnant peptides derived from 469 proteins were identified in free merozoites. Twenty-six proteins were identified as ubiquitylated at all four stages of development, including six of seven histones. Ubiquitylation of many more proteins appeared to be stage specific; for example, seven ubiquitylation sites in 41 protein substrates and 653 ubiquitylation sites in 322 protein substrates were identified only in schizonts and merozoites, respectively (Figure 1). Only six proteins were found to be ubiquitylated exclusively in ring stages, of which three are rhoptry proteins. Our bioinformatics analysis below is extrapolated from these qualitative label free experiments since our efforts to apply quantitative mass spectrometry approaches based on isotope labelling such as TMT or dimethyl labelling [27] were not compatible with the relatively low yield of modified peptides.

### The function and location of selected ubiquitylated proteins

A GO enrichment analysis of the merozoite ubiquitome compared with that of the combined intracellular life cycle stages (rings, trophozoites and schizonts) was performed. Enriched GO terms are depicted in a network representation to visualise dependencies between enriched GO terms as edges between nodes (GO terms). The enrichment map (Figure 1C and 1D, Supplementary Table 4) revealed a large cluster of GO terms associated with protein degradation in the ubiquitomes of both intra- and extra-erythrocytic parasites. The analysis also suggested potential regulatory roles for ubiquitylation in energy metabolism, chromatin and DNA packaging, ribosomes, and vesicles in intra- and extra-cellular life cycle stages. Merozoite-specific clusters of enriched GO terms involved in protein transport, GTP-dependant signalling and the pellicle were observed. A further functional insight into the ubiquitylated merozoite proteins was obtained from a pathway enrichment analysis using the manually curated Malaria Parasite Metabolic Pathways (MPMP) database [28] (Supplementary Figure 2), which suggested a potential crosstalk between ubiquitylation and other post translation modifications such as phosphorylation, myristoylation, and nitrosylation with histone acetylation and methylation as the most enriched term. On the basis of the GO and functional enrichment analyses we focused on some groups of proteins, and three examples will be described here: proteins of the pellicle, exported proteins and histones, as well as ubiquitin itself.

#### Ubiquitylation of pellicle components

The parasite pellicle, a structure composed of the parasite plasma membrane (PM) and the IMC, is important in erythrocyte invasion. The protein components of the pellicle include those attached to the PM inner leaflet, those in the space between the PM and the IMC, such as components of the glideosome, and proteins of the IMC. Eighteen IMC/glideosome components are ubiquitylated, represented by 103 sites (Supplementary Table 5), including myosin A and its light chains and the glideosome associated proteins GAP40 and GAP45, but not GAP50, which is an IMC lumenal protein. There are multiple ubiquitylation sites in several IMC1 proteins, and likely IMC-associated proteins such as PF3D7_0822900 [29] and Phil1 [30] are ubiquitylated; in the case of PF3D7_0822900 25 sites were identified. The vast majority of these sites and substrates were only detected in merozoites (94 sites, 14 proteins).

#### Ubiquitylation of exported proteins

Proteins that are exported beyond the parasite to the infected red blood cell were found to be ubiquitylated (Supplementary Table 6). These proteins can be classified into several groups; for example, there are proteins that are exported from the intracellular parasite and have a PEXEL/HT motif, those transferred from the merozoite at or just after invasion, and proteins that may be targeted to the apicoplast.

Within the data set there are 86 proteins annotated as exported in PlasmoDB (www.PlasmoDb.org), and defined by the presence of an N-terminal signal peptide; the presence of a PEXEL/HT targeting motif [31, 32]; or by reference to the literature, including PEXEL/HT negative exported proteins (PNEPs) [33, 34]. Of these proteins 55 had a signal peptide and of these 10 also contained a PEXEL/HT motif. Of the 31 proteins without a signal peptide, 24 contained a PEXEL/HT motif. Many of these proteins were detected in the ubiquitome at a single stage of parasite development (Supplementary Table 6); whilst 20 exported proteins were detected as ubiquitylated at two or more stages, 34 were only detected in schizonts, 29 were only detected in merozoites and three were only detected in rings. No ubiquitylated exported proteins were detected exclusively in trophozoites. Twenty-two of the proteins annotated as exported that were only ubiquitylated in schizonts contain a PEXEL/HT motif. Exported proteins ubiquitylated in schizonts include many known to be transferred to the host red blood cell (RBC) including KAHRP [35] and REX1 [36] (both with 25 ubiquitylation sites), MESA [37], EMP3 [38], members of the pHIST [39, 40] and PTP families [41, 42], and Maurer’s cleft proteins [43]. In contrast, only five of the exported proteins ubiquitylated only in merozoites contain a PEXEL/HT motif, but 25 of them have a signal peptide. It is possible that this group includes proteins targeted to the apicoplast [44], which has its own ubiquitylation machinery [9], although most of the proteins are not present in the recently described apicoplast proteome [45]. The three exported proteins detected as ubiquitylated only in rings are likely inserted by the invading parasite (but they were not detected in the merozoite ubiquitome).

#### Histone ubiquitylation

There are seven plasmodial histones and they are known to be subjected to a range of posttranslational modifications, including one ubiquitylation site on H2B [46, 47]. We identified between four and five ubiquitylation sites on all histones and ubiquitylation of each histone at all stages except for H2A.Z (PF3D7_0320900) (Supplementary Table 7). There is overlap between the ubiquitylation sites and, for example, methylation and acetylation sites [46] consistent with an involvement of ubiquitylation in the control of gene expression.

#### Polyubiquitylation

Our analytical approach cannot distinguish between the products of monoubiquitylation – the addition of a single ubiquitin to a substrate lysine side chain - and polyubiquitylation – the further ubiquitylation of the attached ubiquitin. Furthermore, the analysis of the peptides derived from polyubiquitylation is potentially confounded by polyubiquitylation of host proteins present in the cell lysates. Since parasite and host ubiquitin differ only by a single residue, the provenance of most ubiquitin-derived diGly-remnant peptides cannot be distinguished except for those in the vicinity of this difference. We identified peptides derived from ubiquitylation at residues 6, 29, 33, 48 and 63 of either host or parasite ubiquitin; ubiquitylation at position 11 from both host and parasite ubiquitin and ubiquitylation at position 27 of only parasite origin (Supplementary Tables 1 and 2). These data are consistent with polyubiquitylation occurring in the parasite, which is also supported by western blotting data obtained using ubiquitin K48- and K63-specific antibodies (see below), but the full extent cannot be assessed by these methodologies.

### Inhibitors of the ubiquitylation pathway inhibit parasite growth and development

The extent and the changes in the pattern of protein ubiquitylation suggested that this modification is important in parasite development and that it would be of interest to test this idea by examining whether or not inhibitors of the ubiquitin pathway affect parasite development.

A number of commercially available inhibitors of E1, E2, E3 and DUB function were tested as inhibitors of parasite growth. As shown in Table 1 and Supplementary Figure 3 some of these compounds were effective inhibitors, although their mode of action in the parasite has not been validated. In particular, the EC_50_ for MLN7243 (an E1 inhibitor) and SMER3 (an E3 inhibitor) were in the sub-micromolar range. Given that MLN7243 is an inhibitor of E1 (UBA1, ubiquitin activating enzyme) and that ubiquitin activation is the first step in the ubiquitylation pathway, it was of interest to investigate the activity of this compound in more detail.

**Table 1.** Potency of various ubiquitylation enzyme inhibitors against *P. falciparum in vitro*.

|  |  |
| --- | --- |
| **Target** | **EC<sub>50</sub> (μM)** |
| **E1 (ubiquitin activation)** |  |
| MLN7243 | 0.4 |
| PYR41 | 28 |
| PYZD4409 | 133 |
| **E2 (ubiquitin conjugation)** |  |
| BAY 11-7082 | 21 |
| NSC697923 | 32 |
| **E3 (ubiquitin ligation)** |  |
| Heclin | 38 |
| Nutilin-3A | 2.3 |
| SMER3 | 0.8 |
| **DUB (ubiquitin removal)** |  |
| PR619 | 2.3 |
| Inhibitors of ubiquitin addition (E1, E2 and E3 activities) or removal (DUB activity) were tested and the EC<sub>50</sub> calculated from the graphs in Supplementary Figure 3. |  |

### MLN7243 inhibits parasite development at the transition from schizonts to merozoites

We examined the effect of MLN7243 on intra-erythrocyte development, merozoite release and re-invasion to form new ring stages, as well as on protein ubiquitylation in the parasite (Figure 2). Synchronised parasite populations were treated with either 1.6 µM MLN7243 or DMSO immediately after invasion. Parasite growth and multiplication were examined by FACS analysis, and parasite morphology was assessed using light microscopy of Giemsa stained slides. In DMSO-treated cultures parasites transitioned from ring stages with a single nucleus at 3 h post-invasion (hpi) to multinucleate schizonts at 44 hpi and then, following invasion, back to ring stages (Figure 2A, B) with an increase in parasitaemia (Figure 2C). In contrast, MLN7243-treated parasites progressed to the schizont stage (Figure 2A and B) but did not release merozoites or reinvade to form new ring stages (Figure 2A, B, and C). Washout of the drug at the schizont stage did not restore growth. Analysis of schizonts from DMSO or MLN7243-treated cultures by western blot with ubiquitin-specific antibodies revealed that in the presence of MLN7243 there was a marked reduction in detectable ubiquitylation, including the K48- and K63-specific modifications of ubiquitin that are characteristic of polyubiquitylation (Figure 2D). These results indicate that the inhibitor is acting to prevent the maturation of the intracellular schizont and therefore inhibits the release of viable merozoites, and that its action is irreversible.

**Figure 2.**
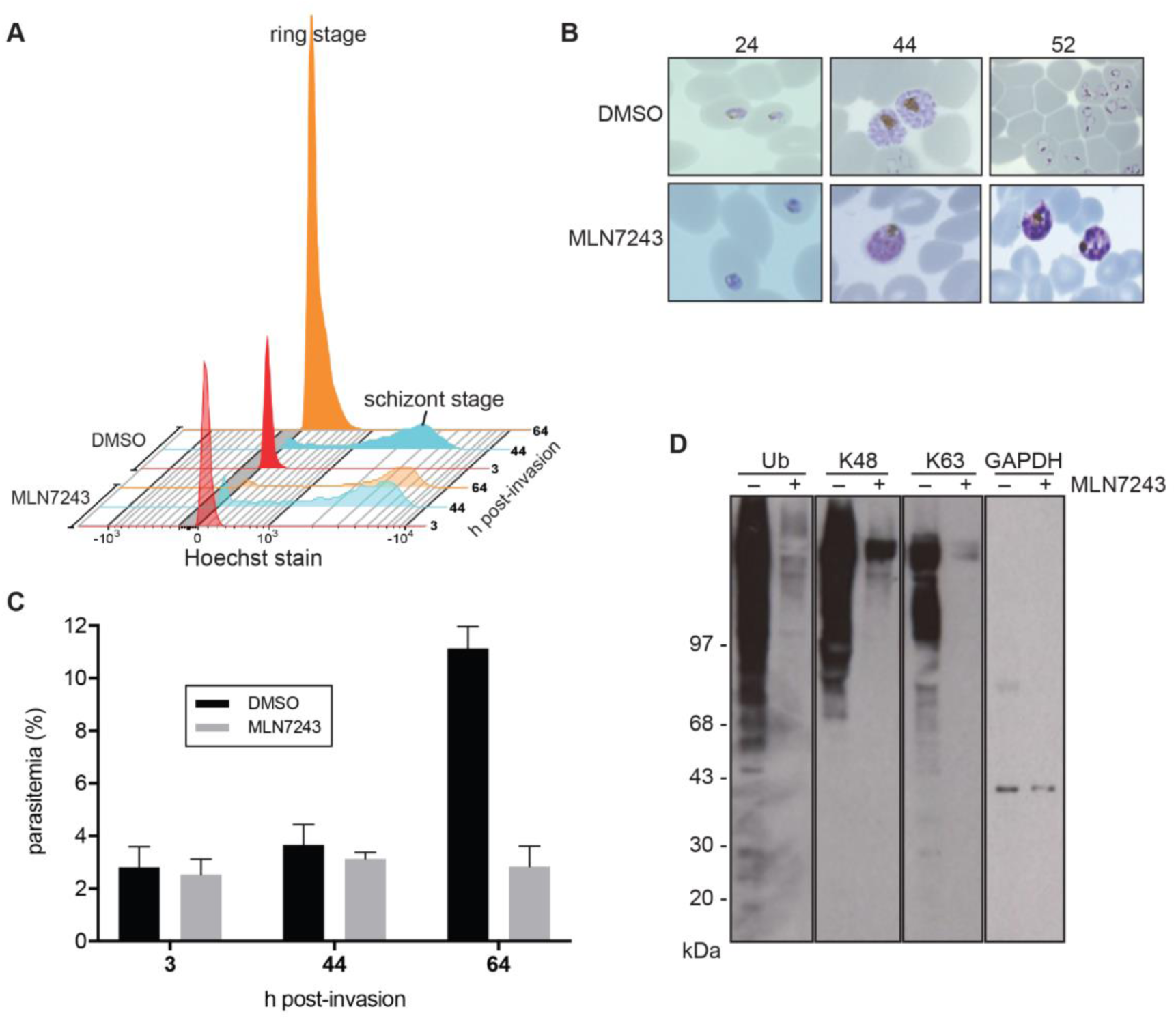
Treatment with MLN7243 arrests parasites at the schizont stage and blocks protein ubiquitylation. (A) Parasites were treated with either MLN7243 or DMSO carrier, stained with Hoechst dye and analysed by FACS at three time points: 3, 44, and 64 h post invasion, shown in red, blue and orange respectively. Haploid ring stages transition to multinucleate schizonts (shown at 44 h) and in the control DMSO-treated cultures, merozoite reinvasion produces new haploid intraerythrocytic parasites (shown at 64 h). The MLN7243-treated parasites do not release merozoites or form new haploid intraerythrocytic parasites. (B) Microscopy of Giemsa-stained parasites at 24, 44 and 52 h post invasion, showing that both cultures develop into schizonts but by 52 h control parasites have formed new ring stages and the MLN7243-treated parasites remain as schizonts. (C) The percentage of parasitaemia in cultures at 3, 44 and 64 h post infection measured by FACS analysis. During intracellular development there is no change in parasitaemia, but by 64 h the parasitaemia has increased only in the control (DMSO-treated) culture. (D) Protein ubiquitylation in schizonts either treated (+) or untreated (-) with MLN7243 during development from ring stages. Western blots were probed with a ubiquitin-specific antibody or antibodies specific for K48- and K63-linked polyubiquitylation. Antibodies to parasite GAPDH were used as a loading control.

### Structural and Biochemical Studies: MLN7243 inhibits recombinant PfUBA1 encoded by PF3D7_1225800

Although it is likely that the observed effect of MLN7243 is via inhibition of UBA1, and manifests at the stage of the cycle when there is a large increase in protein ubiquitylation, we wished to establish that MLN7243 indeed inhibits parasite UBA1. Therefore, we undertook modelling and biochemical studies to investigate this. MLN7243 is an adenosyl sulphamate that forms a UBA1-catalysed covalent adduct with ubiquitin following ATP-dependent activation of ubiquitin, and inhibits both the human and yeast UBA1 (E1) enzymes (HsUBA1 and ScUBA1). However, it has not been established that it targets UBA1 in the malaria parasite. PF3D7_1225800 has been identified as the gene likely to code for the ubiquitin E1 [8], and therefore, we wished to establish that the PF3D7_1225800 product is PfUBA1 and that it can be inhibited by MLN7243 *in vitro*.

Support for MLN7243 targeting of the putative parasite UBA1 was provided by a homology model of the enzyme (Figure 3A) based on the structure of the yeast UBA1-ubiquitin complex with MLN7243 (Figure 3B) covalently linked to the ubiquitin C-terminus [19]. There is high sequence conservation between yeast and *P. falciparum* UBA1 (overall sequence identity of 39.6%), in particular in the active adenylation domain (sequence identity of 78.6% for residues within 5 Ångstrom (Å) of MLN7243), and therefore the model was expected to reproduce many aspects of the *P. falciparum* UBA1-Ub-MLN7243 structure. The PfUBA1-Ub and ScUBA1-Ub complexes are quite similar as reflected in a root mean square (rms) deviation of 2 Å for 927 out of 1006 residues present in ScUBA1. More importantly, PfUBA1 can interact with MLN7243 in a highly similar fashion to the yeast enzyme. This is reflected by the fact that within a radius of 5 Å from MLN7243 the rms deviation between the model of the PfUBA1-ubiquitin-MLN7243 complex and the ScUBA1-ubiquitin-MLN7243 structure drops to 0.76 Å. Surface representations of the ATP binding pockets of human (PDB entry 6DC6), yeast (PDB entry 5L6J) and the PfUBA1 homology model (Figure 3C) indicate that the regions occupied by Mg^2+^-ATP are nearly identical. The only difference is the formation of a weak salt bridge between Lys604 and Asp629 in PfUBA1 where these two residues are within 5.4 Å of each other, in contrast to ScUba1 and HsUBA1; in the latter the Lys is replaced by Arg, and the minimal distance between these oppositely charged residues is in excess of 7 Å. This apparently large discrepancy in surface representation is due to a different side chain rotamer of the positively charged residue in yeast and human UBA1 (χ1 of -132° and -180°, respectively) compared with PfUBA1 (χ1=-80°). Almost all of the residues contacting MLN7243 in ScUba1 are conserved in PfUBA1 and are expected to engage in similar interactions with MLN7243. Notable exceptions are Ala607 of PfUBA1 (corresponding to Pro554 in HsUBA1 and Pro522 in ScUBA1), Gln632 (Asp579 in HsUBA1 and Asp547 in ScUba1) and Asn639 in PfUBA1 (Arg586 in HsUBA1 and Arg554 in ScUBA1) as shown in Figure 3D. All three residues are located at the distal end of the ATP-binding pocket, away from the covalent linkage to ubiquitin, and their different side chains are responsible for altering the shape of the binding pocket in this region. As a consequence of these substitutions the CF_3_-group of MLN7242 in the homology model is rotated by ∼90° around the S-C bond to better match the PfUba1 surface.

**Figure 3.**
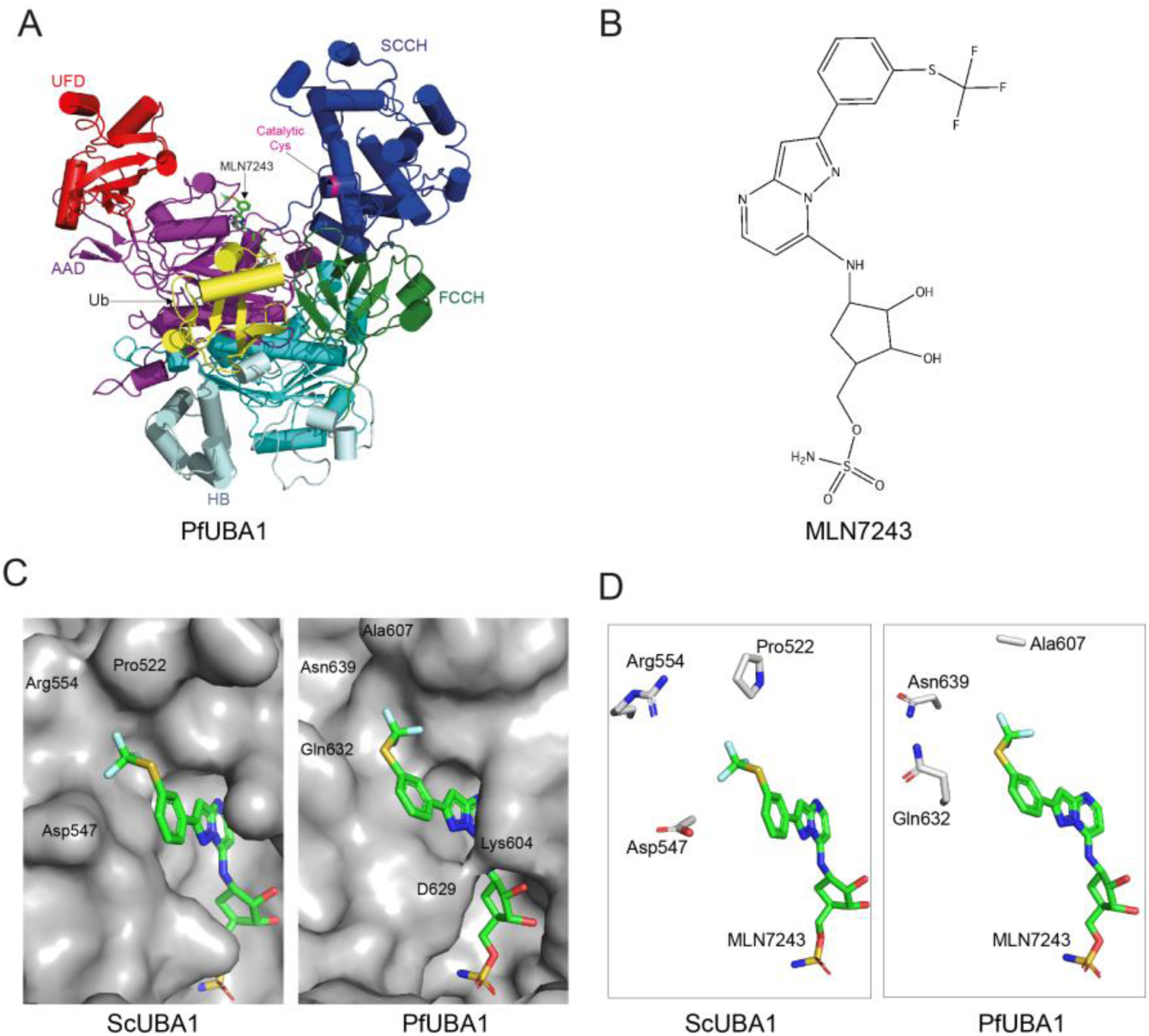
Homology model of PfUBA1 in complex with ubiquitin and MLN7243. (A) Overall structure of the ternary PfUBA1-ubiquitin-MLN7243 complex with α-helices drawn as cylinders and β-strands as arrows. Ubiquitin (Ub) is shown in yellow, PfUBA1 is coloured according to its domain architecture (AAD [purple] and IAD [teal]: active and inactive adenylation domains, HB [cyan]: helix bundle, FCCH [green] and SCCH [blue]: first and second catalytic cysteine half domains, UFD [red]: ubiquitin fold domain) and MLN7243 is shown in stick representation. (B) Chemical structure of MLN7243 (TAK-243). (C) Surface representations of ScUBA1 (left) and PfUBA1 (right) in the vicinity of MLN7243 which is coloured according to atom type with C-atoms in green, O-atoms in red, N-atoms in blue, S-atoms in yellow and F-atoms in light cyan. The locations of residues discussed in the text are indicated. The covalent link between MLN7243 and ubiquitin via the terminal N-atom is not shown. (D) Differences in the residues surrounding the CF_3_-group of MLN7243 are highlighted. HsUBA1 features the same residues indicated for ScUBA1, namely Pro554, Asp579 and Arg586.

We purified the protein from a codon-optimized gene encoding PF3D7_1225800 (Supplementary Figure 4) expressed in insect cells and examined its activity using ubiquitin-E1 thioesterification and ubiquitin-E1 and E2 trans-thioesterification assays. The purified protein (Supplementary Figure 5) appeared to be folded as judged by nano-differential scanning fluorimetry and light scattering with a melting temperature of approximately 45°C (Supplementary Figure 5). To establish whether the protein was enzymatically active, it was incubated in the presence of ubiquitin labelled with a fluorescent dye, and the formation of a UBA1∼Ub thioester with the putative UBA1 catalytic site cysteine was monitored. In the absence of ATP no thioester formation occurred (Figure 4A), but in the presence of ATP, PfUBA1 was conjugated with the fluorescently labelled ubiquitin, indicating that the protein was catalytically active. Furthermore, the enzyme was able to transfer ubiquitin to four different human E2s (UBCH5, UBCH7, CDC34 and UBC13) in a manner identical to that of the murine enzyme (Mm UBA 1, Supplementary Figure 5), confirming its ubiquitin activation activity. Increasing concentrations of MLN7243 in the Ub∼UBA1 thioesterification assay reduced the amount of bound ubiquitin, allowing an IC_50_ of 1.4 ± 0.2 µM to be calculated (Figure 4B and 4C). In contrast, MLN4924, an adenosyl sulphamate inhibitor of Nedd8 E1 [48] showed a much higher IC_50_ (47 ± 5 µM) for the inhibition of thioester formation (Supplementary Figure 5).

**Figure 4.**
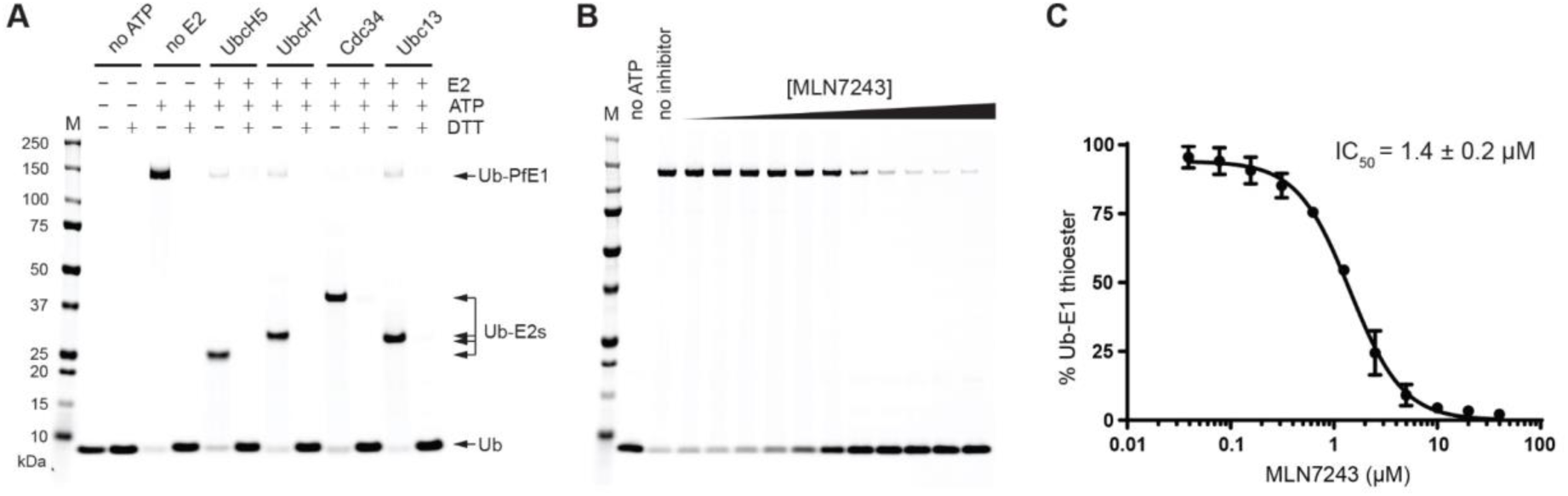
Thioesterification of PfUBA1 E1 using fluorescently labelled ubiquitin (Ub) and the transthioesterification of four distinct E2s *in vitro*. (A) In the presence of Pf UBA1, thioesterification of the E1 by Ub occurs in the absence of E2 and in the presence of four distinct E2s (UBCH5, UBCH7, CDC34 and UBC13) further transthioesterification to the E2 occurs. No thioester formation is observed in the absence of ATP or after treatment with DTT. (B) Increasing concentrations of MLN7243 reduce covalent addition of ubiquitin to PfUBA1, allowing (C) the calculation of its IC_50_. Data are shown as mean ± SD from triplicates. Expression, purification and further characterization of the recombinant PfUBA1 and additional control experiments are provided in Supplementary Figures S4 and S5.

These data indicate that PF3D7_1225800 encodes an active ubiquitin activating enzyme (PfUBA1) that can be inhibited by MLN7243 and is therefore the likely target of this inhibitor in the parasite.

### Both an inducible knockout of PF3D7_1225800 and treatment with MLN7243 stop schizont development

Since adenosyl sulphamates may also inhibit other processes, such as Neddylation, we wished to confirm that MLN7243 inhibition of UBA1 caused the parasite death that we observed. We generated parasite cell lines with an inducible genetic functional knock out of the *uba1* gene PF3D7_1225800 to examine whether gene loss recapitulates the inhibitor-induced phenotype.

An inducible functional knockout of PF3D7_1225800 was generated using CRISPR-Cas9 technology. The genetic manipulation was carried out in the II-3 parasite line, which contains the rapamycin-dependent di-Cre recombinase inserted in the genome [24]. PF3D7_1225800 contains two introns, with one located immediately upstream of the 3’ region that encodes the C-terminal ubiquitin fold domain (UFD). We replaced the second endogenous intron with a *P. falciparum sera2* intron containing a loxP site (loxPint) [23], added a GFP coding sequence to the end of the gene and placed a second loxP site in the 3’UTR, immediately followed by sequence coding for a triple HA tag. Upon inducible expression of Cre recombinase a truncation of the modified gene occurs, resulting in an inactive protein product that will contain a triple HA-tag at its C-terminus. A schematic of the strategy to manipulate PF3D7_1225800 is shown in Figure 5A. Following transfection and selection, three clonal parasite lines were chosen for further characterization. Each of the three lines was generated using the same repair plasmid but a different guide RNA, so they should be genetically identical. The GFP-tagged protein was expressed in the absence of rapamycin, and the addition of the tag had no detectable effect on parasite growth. Synchronised parasite populations were treated with rapamycin approximately four hours after red blood cell invasion and then analysed for Cre-mediated sequence deletion by PCR and loss of GFP expression by western blot during the schizont stage, approximately 40 h later. The PCR analysis indicated that modification of PF3D7_1225800 had been successful, and that rapamycin treatment resulted in excision of the sequence between the loxP sites (Figure 5B). Western blot analysis showed loss of the GFP signal upon rapamycin treatment (Figure 5C), consistent with deletion of the C-terminal ubiquitin fold domain and the GFP tag. However, we saw no appearance of a truncated, HA-tagged protein after rapamycin treatment (data not shown), which suggests that the truncated protein is unstable and rapidly degraded.

**Figure 5.**
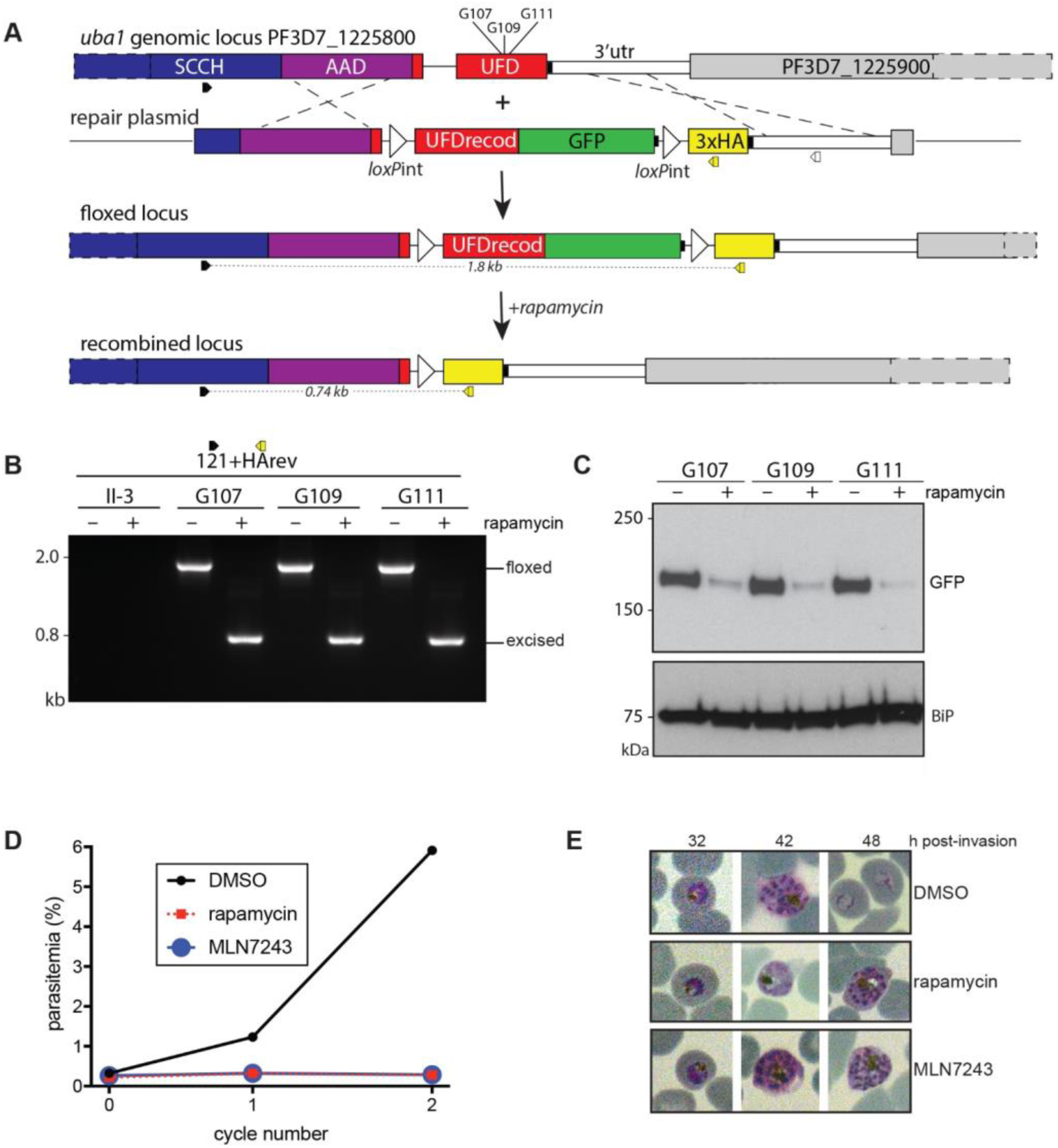
Generation of a parasite line with a rapamycin-inducible *uba1* gene knockout and its response to treatment with either rapamycin, MLN7243, or DMSO. (A) CRISPR Cas9 was used to modify the 3’ end of the *uba1* gene in the II-3 parasite line, inserting loxPint in place of the last intron, a recodonised coding sequence fused to GFP, a second loxPint sequence and sequence coding for a triple HA tag. The parasite has a floxed gene from which a UBA1-GFP fusion protein is expressed. In the presence of rapamycin the floxed sequence is excised to produce an inactive truncated UBA1 protein. (B) Diagnostic PCR with primers designed to detect successful integration of the floxed sequence and its excision in the presence of rapamycin. DNA from the parental (II-3) and three cloned recombinant lines (G107, G109 and G111), following growth with or without rapamycin, was amplified using the primers 121 and HArev. Amplified bands of 1.8 kb and 0.74 kb were expected from the floxed and excised (+rapamycin) locus in the progeny parasites and no product was expected from the parental II-3. (C) Western blot with anti-GFP antibodies probed against schizont extracts of parasites (clones G107, G109 and G111) with the floxed and excised (+ rapamycin) *uba1* locus. Reactivity of antibodies to BiP(Binding immunoglobulin Protein, GRP-78) was used as a loading control. (D) Synchronous G109 parasites were treated at the early ring-stage with either DMSO (control), rapamycin, or MLN7243 and the parasitaemia in each culture was monitored for two growth cycles. Genetic truncation or inhibition of UBA1 blocked parasite growth and reinvasion. (E) Images of Giemsa-stained parasites treated with either DMSO, rapamycin or MLN7243. Parasites in all conditions develop into schizonts but, whereas the control parasites develop further into ring stages, drug treated parasites do not proceed beyond the schizont stage.

To directly compare the parasite phenotype resulting from the inducible functional knockout of PF3D7_1225800 and treatment with MLN7243, synchronised floxed *uba1* G109 parasites were treated with either rapamycin (to truncate the gene), MLN7243 (to inhibit the enzyme), or DMSO (control) immediately after invasion, and parasite development examined by FACS and light microscopy. The DMSO-treated parasites developed normally, showing an increase in parasitaemia (Figure 5D) and formation of ring stages (Figure 5E), whereas both the rapamycin and MLN7243 treated parasites stalled during schizont development and failed to produce any merozoites so that there was no associated increase in parasitaemia. These results show that the phenotype resulting from the genetic knock out of PF3D7_1225800 is the same as that resulting from treatment of the parasite with MLN7243, consistent with the on-target activity of this inhibitor against PfUBA1.

## Discussion

We describe a parasite asexual blood stage ubiquitome that is much larger than was known previously. Although the majority of the di-Gly remnant peptides were likely derived from ubiquitylated proteins, some peptides may have been derived from Neddylated proteins such as the Cullin subunits of SCF ubiquitin ligases, but the number of these is expected to be small [49]. In previous studies, potentially ubiquitylated proteins were identified by binding to TUBES resin or a ubiquitin-specific antibody, and then identified by mass spectrometry. These methods do not identify the site of modification that provides the strongest evidence of ubiquitylation. Of the 106 proteins identified by Mata-Cantero and colleagues [12], 56 (52%) were also detected in our study. Of these 106 proteins, twelve were also identified in the earlier Ponts study [11] and of these, ten were detected in our study. It is possible that these earlier approaches selectively enriched proteins that are highly polyubiquitylated, whereas our approach also detected proteins with one or more ubiquitylation sites independent of the length of the attached ubiquitin chain. Mata-Cantero and co-workers specifically detected 17 ubiquitylation sites in five parasite proteins and three of these proteins were also identified in our analysis. The earlier studies did not examine the extracellular merozoite, which we now show is where most ubiquitylation is present. It is clear that our detection of 1464 ubiquitylation sites in 546 parasite proteins represents a considerable increase in the size of the known parasite ubiquitome.

The ubiquitomes of three intracellular stages (ring, trophozoite and schizont) and extracellular merozoites were examined. The ubiquitylation sites were derived from both parasite and host erythrocyte proteins; it is known that erythrocytes contain a residual ubiquitin machinery [50]. We used saponin treatment to remove soluble erythrocyte proteins, but this did not remove all erythrocyte components. As a consequence of the human protein contamination and the near identity of the parasite and human ubiquitin sequences it was impossible to determine the extent of polyubiquitylation that occurs in the parasite. However, the reactivity of parasite proteins with antibodies specific for the Lys47 and Lys63 linkages in polyubiquitin, which was sensitive to parasite treatment with MLN7243, suggests that polyubiquitylation in the parasite is extensive. The sites in parasite ubiquitin at residues 11 and 27 could be positively identified due to the sequence difference between human and Pf ubiquitin, however, it is likely that the parasite ubiquitylates ubiquitin at all seven lysine residues, since corresponding diGly-remnant peptides were identified in the merozoite samples.

We classified the substrates to identify those that are enriched in the ubiquitome and focused on four groups. It is of interest that several components of the IMC, the apical organelles and the glideosome motor are ubiquitylated in the merozoite, consistent with their destruction following invasion. For example, of the 18 proteins associated with the pellicle 14 are ubiquitylated only in merozoites. A number of proteins known to be exported to the host RBC are ubiquitylated, particularly in schizonts. It is possible that these proteins have not been secreted, but modified and retained in the parasite cytoplasm, for example via an ERAD (endoplasmic reticulum associated protein degradation) system that targets misfolded secreted proteins and moves them to the cytosol for subsequent proteasomal degradation. Alternatively, these proteins may have been ubiquitylated following their arrival in the host cell cytosol, implying that the RBC ubiquitylation system is able to modify exported proteins of the parasite. In the case of the KAHRP, which is located in knobs formed in trophozoite-infected erythrocytes, it may be significant that the protein was only found to be ubiquitylated in schizonts despite the fact that the gene is transcribed from the ring stage [35]. It has been suggested previously [12] that the erythrocyte ubiquitylation system is modulated as a result of infection. If the host machinery is responsible, then host E3 ligases can recognise parasite proteins. However, it is unclear whether this modification has functional significance for the parasite, for example in cytoadhesion, or in parasite egress. The ubiquitylation of histones throughout the cell cycle was found to be extensive and suggests that this modification needs to be considered in investigations of chromatin-dependent gene regulation. Ubiquitylation of parasite histones at sites that are not present or modified in human histones may, in part, explain why antibodies raised to parasite histones do not cross react with human histones [51]. The identification of ubiquitylation sites in many other parasite proteins opens up the possibility to examine the importance of this modification in their location and function.

From a range of inhibitors of ubiquitylation in other organisms, we showed that some inhibited parasite growth *in vitro,* suggesting that ubiquitylation and deubiquitylation may be attractive targets for the development of novel anti-malarial drugs. Such inhibitors can provide insights into novel targets for anti-malarial therapy, provided that their mode of action and specificity are conserved. Of the compounds tested here, the E1 inhibitor MLN7243 was the most potent, although strong anti-parasite inhibition was also shown by the E3 ubiquitin ligase inhibitors SMER3 and Nutlin-3 as well as the DUB inhibitor PR619. Interestingly Jain and colleagues [21] showed that certain known E3 inhibitors can inhibit malaria parasite growth in vitro, including the two compounds, Nutlin-3 and SMER3 that we tested. For any inhibitor it needs to be established that the mode of action is on-target rather than via an alternative mechanism. We chose to validate the mode of action of the most potent of the available inhibitors, MLN7243, using *in vitro* biochemical and *in vivo* genetic approaches. Treatment of parasites with MLN7243 or inducible inactivation of *Pfuba1* both resulted in a block of parasite development at the transition from intracellular schizont to extracellular merozoite, which corresponds to the point where there is a large increase in ubiquitylation in the cell. Although some protein ubiquitylation occurs prior to this point in the life cycle, it is possible that there is sufficient activated ubiquitin present in the cell to fulfil requirements prior to the increase necessary for merozoite maturation, alternatively specific ubiquitylation pathways late in schizont development, such as a role in egress, may be preferentially affected. There is another ubiquitin E1 encoded in the genome (PF3D7_1333200), which is targeted to the apicoplast, and it is not clear whether or not this could be a source of activated ubiquitin in the cytoplasm or whether it is also targeted by MLN7243. However, the genetic evidence suggests that an E1 other than Pf3D7_1225000 is not responsible for the observed phenotype.

MLN7243 is a member of the adenosyl sulphamate class of compounds that target E1 enzymes responsible for ATP-dependent activation [52], including MLN4943 that targets the E1 involved in Neddylation and which may also be important in the parasite [53]. New and potentially more active UBA1 inhibitors will need to show enhanced potency and good selectivity over the host enzyme. The homology model of the ternary PfUBA1-ubiquitin-MLN7243 complex suggests that design of related adenosyl sulphamates with high specificity for PfUBA1 over HsUBA1 should be possible, due to the differences in the residues that define the properties of the inhibitor binding pocket at its distal end. Arg586 of HsUBA1 was previously suggested as a residue that could mediate specificity for the development of inhibitors of HsUBA1 over other E1 enzymes present in humans [19], and its replacement with the shorter Asn639 side chain in PfUBA1 suggests that inhibitors with extensions in the distal direction may specifically attack PfUBA1. The evolution of parasite resistance to such inhibitors is also an important concern that should be address early in the drug discovery process [25]. A previous study showed that upon MLN7243 treatment HsUBA1 Ala580, which has been designated as the gatekeeper residue, is replaced with a serine [54], thereby providing drug resistance. PfUBA1 has an alanine at this position (Ala633) and hence drug resistance to adenosyl sulphamates might develop in *P. falciparum* in the same way. The yeast enzyme is structurally more similar to that of the parasite than the human, opening opportunities for the easier development of new antimalarial compounds. New candidate compounds developed in this area will need an ADMET (absorption, distribution, metabolism, excretion, toxicity) profile, and other characteristics [55] appropriate for an antimalarial drug.

Protein ubiquitylation plays many key roles in eukaryotic cells. The best understood is its role in targeting proteins to the proteasome for degradation, although other critical roles include sorting proteins to subcellular locations. The proteasome has a central role in parasite biology and specific proteasome inhibitors kill malarial parasites, thereby identifying inhibitors of the ubiquitin-proteasome system as possible anti-parasitic reagents [56–58]. Ubiquitylation is important in mediating the effects of artemisinin. Parasite treatment with artemisinin results in activation of the unfolded protein response and the accumulation of polyubiquitylated substrates, compromising proteasomal function [16, 59, 60], and the proteostatic dysregulation of parasite phosphatidylinositol 3-kinase (PI3K) [61]. Amino acid substitutions in the K13 Kelch protein are associated with artemisinin resistance, and it has been proposed that K13 is a ubiquitin E3 ligase adaptor for PI3K [61]. K13 and PI3K associate with the parasite ER and potentially involved in quality control of exported and cytoplasmic proteins [62, 63]. Interestingly an E1 inhibitor antagonises dihydroartemisinin activity [16] and therefore such compounds would not be useful in combination with artemisinin, but DUB inhibitors such as PR619 may be useful in this regard.

We propose the following model for one important aspect of ubiquitylation in parasite biology at the transitions between intracellular and extracellular parasites, where there are large differences in the extent of ubiquitylation. The enhanced ubiquitylation of proteins in the extracellular merozoite is consistent with a role in their proteasomal degradation following invasion, which allows recycling of these resources at a time when they have no further function and other nutrient sources may not exist. Normally the intracellular parasite acquires nutrients required for growth from the host cell by digestion of host proteins in the food vacuole, an acidic compartment that is elaborated anew each intracellular cycle and then discarded at parasite egress as the residual body. The merozoite has no food vacuole and no lysosomal compartment and therefore, in the absence of lysosomal or vacuolar mechanisms for protein catabolism, it is likely that after invasion the new ring stage parasite relies on the ubiquitin-proteasome system for protein turnover to supply amino acids until a new food vacuole is functional. Consistent with this idea, the intracellular structures such as the IMC and secretory organelles are disassembled and disappear, and it is likely that the components are recycled. This model predicts that proteasomal function (and therefore the effect of specific inhibitors) is particularly important in ring stage parasites, and it is interesting that proteasome inhibitors [64] and artemisinin [65] are important for targeting this stage of parasite development.

## Materials and Methods

### Parasitology

#### Parasite culture and preparation of ring, trophozoite, schizont and merozoite stages for ubiquitylation analysis

*P. falciparum* 3D7 parasites were cultured at 37°C *in vitro* in plastic tissue culture flasks (Nunc^TM^) at between 1 and 10% parasitaemia and 1 to 4% haematocrit, as described [66]. The culture medium was comprised of RPMI 1640 supplemented with 1% (w/v) Albumax, and was replaced every 2 - 3 days at which time the cultures were also gassed (5% CO_2_, 5% O_2_, 90% N_2_) and the flasks sealed. Parasite populations were synchronized by collection of schizonts following centrifugation on a 70% Percoll cushion. The schizonts were returned to culture for 2 h to allow invasion of merozoites into fresh red blood cells and then 5% D-sorbitol was added for 10 min to kill all parasites except the young ring stage [67]. After a further centrifugation at 1800 g for 3 min, the parasitized red blood cells were resuspended in fresh medium and the flasks gassed. These synchronous cultures of parasites were used to prepare 500 µl of ring-, 500 µl of trophozoite- and 1000 µl of schizont-stage parasites. Parasites were lysed in 0.15% (w/v) saponin in PBS to remove haemoglobin, and washed in PBS prior to snap-freezing in dry ice and storage at -20 °C. Merozoites were purified using a magnetic column as described previously [66, 68]. Approximately 600-700 µl of merozoites that had been washed in PBS were used.

#### Parasite drug treatment and SYBR green assay

To determine the EC_50_ of compounds, we measured parasite growth over two cycles using a fluorescence-based SYBR Green assay performed as described previously [69]. Final compound concentrations were established by serial twofold dilutions with an upper concentration of 10 µM for MLN7243; 50 µM for BAY 11-7082, SMER3, Nutlin-3A, and PR61; and 100 µM for PYR41, PYZD4409, NSC697923, and Heclin. The final DMSO concentration was maintained at 0.05% in all cases. Aliquots of 100 µl *P. falciparum* culture 24 h after erythrocyte invasion were placed into 96-well culture dishes. Cells at a starting parasitaemia of 0.2% were incubated for 96 h with inhibitors dissolved in DMSO. After incubation, cells were lysed by the addition of 25 µl buffer (20 mM Tris-HCl, pH 8.0, 2 mM EDTA, 1.6% Triton X100, 0.16% saponin, 5x SYBR Green I [Life Technologies]) and incubated in the dark for 2 h. Parasite growth was determined by measuring the fluorescence of the samples using a FLUOStar Omega plate reader (BMG Labtech) with excitation and emission filters of 485 nm and 520 nm, respectively [69]. EC_50_ values were calculated from a four-parameter logistical fit of the data using the Prism software (GraphPad Software, Inc.).

### Genetic modification of the parasite

#### Parasite transfection

*P. falciparum* schizont stage parasites were transfected with a 4D-Nucleofector instrument using a P3 Primary Cell kit (Lonza). One day after transfection, drug selection was commenced with 2.5 nM WR99210. Drug selection was applied for four days then removed.

#### Inducible truncation of the uba1 gene

A conditional deletion of the last exon of *uba1* was generated using CRISPR-Cas9 to target nuclease activity to the *uba1* locus in a parasite line expressing a rapamycin-inducible diCre recombinase [24]. Guide RNAs were designed using the Protospacer software (www.protospacer.com) [70] that would target Cas9 to introduce a double-stranded break at nucleotide 4576, 4674, or 4787 in the penultimate exon of the uba1 gene. Pairs of complementary oligonucleotides, 107for/rev, 109for/rev, or 111for/rev were phosphorylated with T4 polynucleotide kinase, annealed, and ligated into pDC2—Cas9-hDHFRyFCU digested with BbsI as described in [24]. A synthetic DNA construct containing 500 bp of the *uba1* gene upstream of its second intron, followed by a *sera2* intron containing a *loxP* site (described in [23], and the recodonized final exon of *uba1* was generated in the pMK vector (Life Technologies). The sequence of the final exon was altered so that the next best codon based on *P. falciparum* codon usage (plasmodb.org) was used to encode the relevant amino acid. Another synthetic DNA fragment containing GFP, a *sera2:loxP* intron, and a triple HA epitope tag was introduced into this plasmid, using AvrII/ SacII restriction sites at its 5’ and 3’ ends, respectively. All synthetic DNA was ordered from GeneArt (Life Technologies). Finally, a DNA fragment comprising the 3’UTR (451 bp) of *uba1* plus 75 bp of the downstream gene was generated by PCR using primers 113 & 114 and cloned into the repair plasmid using SacII/HindIII restriction sites. The repair plasmid (50 µg) was linearized by digestion with BglII, ethanol precipitated with 20 µg guide plasmid DNA, and redissolved in 10 µl sterile TE (10 mM TrisHCl pH 8.0, 1 mM EDTA pH 8.0). Transfection was carried out as described above. Following rapamycin-mediated dimerisation of Cre recombinase the floxed sequence is removed, truncating the protein by removal of the ubiquitin fold domain and GFP sequences and replacing them with the HA tag. Three independent clones with the correct integration, generated using different guide RNAs (G107, G109 and G111), were selected for further analysis. Correct integration of the modified gene was confirmed by PCR. The clones were screened by PCR using the primers 121 and 116 (to generate products of 1129 bp from II-3 [WT] and 2013 bp from the floxed gene, respectively), primers 121 and GFPrev (to generate no product from II-3, and a 949 bp product from the floxed gene), and primers 121 and HArev (to generate no product from II-3 and an 1840 bp product from the floxed gene). With primers 121 and HArev, a 742bp product was expected from the modified locus following Cre-mediated recombination. (Figure 5). The sequences of all oligonucleotides used in this study are displayed in Supplementary Table 8.

### Analysis of parasite development and protein expression

#### Analysis of parasite development by microscopy

For light microscopy, parasite smears were prepared on glass slides and stained with Giemsa’s reagent prior to examination of parasite morphology with a Zeiss Axioplan 2 microscope. Parasitaemia was determined from the number of infected and uninfected erythrocytes and parasite morphology at different stages of development and following drug treatment was also noted.

#### FACS-based assay of parasite development

Synchronised parasite populations were cultured with or without inhibitor. The cells were then fixed in 2% paraformaldehyde, 0.2% glutaraldehyde and parasite DNA stained for 10 min with 2 µg/ml Hoechst 33342 dye. Cells were examined using a BD LSRFortessa X-20 flow cytometer and an argon laser tuned for UV (355 nm), with fluorescence at 465nm; Parasites with one nucleus can be clearly distinguished from those with 2 or more nuclei and were sorted by DNA content [25].

#### Parasite growth assays

The synchronized II-3 and Δuba1 TS cloned lines (at 0.2 or 2.0% parasitaemia, 3% haematocrit) were treated at the early ring stage with DMSO, MLN7243 (3.2 µM final) or rapamycin (100 nM final) and samples taken periodically to assess the consequence of drug addition, using Giemsa staining to assess parasite morphology and FACS analysis to assess growth/reinvasion.

#### Western blotting of parasite lysates

Schizonts were purified by centrifugation over a 63% Percoll cushion, and the erythrocyte membrane was lysed by resuspension in 0.15% saponin in PBS, followed by centrifugation at 5,000 g for 5 min. The resulting cell pellet was dissolved in 10 volumes of lysis buffer (150 mM NaCl containing 1% Triton X-100, 0.1% SDS, 1µl/ml benzonase (Roche), and 1 x complete protease inhibitors (Roche). Samples were incubated on ice for 10 min followed by centrifugation at 17,000 g for 30 min. The supernatant was recovered and its protein content quantified using a BCA Protein Assay (Pierce). Seven µg total protein was resolved on a 3 to 8% Tris-acetate PAGE gel (Invitrogen), transferred to nitrocellulose, and probed with specific antibodies. Protein ubiquitylation was detected with antibodies to ubiquitin (mouse mAb FK2, Merck) and K48 and K63 specific linkages (Millipore rabbit mAbs 05-1307 and 05-1308, respectively). UBA1 protein expression was monitored by western blotting using antibodies to GFP (Roche). Anti-BiP and anti-GAPDH antibodies were used to detect loading control proteins.

### Proteome analysis

#### Parasite lysis and protein digestion

To prevent proteolysis of the ubiquitin linkage to protein substrates, we prepared parasite lysates in a solution containing 9 M urea to denature proteases. These lysates were treated with LysC and trypsin proteases (Promega) to cleave proteins into peptides at arginine and unmodified lysine residues. This treatment also digests ubiquitin attached to lysine sidechains leaving a residual Gly-Gly-remnant peptide. A washed parasite pellet (∼200 µl) was re-suspended in 5 ml of 9 M urea, 20mM HEPES, pH 7.8, supplemented with 500 units of benzonase and sonicated to reduce viscosity (3 mm probe, 50% amplitude, 3 x 15 s bursts, on ice). Between 5 - 10 mg of protein per sample were used as estimated by Bradford protein assay. Lysates were reduced with 10 mM dithiothreitol (DTT) (Sigma) for 30 min at room temperature, followed by alkylation with 20 mM chloroacetamide (Sigma) for 30 min at room temperature in the dark. Lysates were digested initially with LysC (Promega) for 2 h at 37⁰C. The lysates were then diluted with 100 mM ammonium bicarbonate, 5% acetonitrile to a final urea concentration of less than 2 M. The samples were digested at a 1:100 enzyme to protein ratio (w/w) with trypsin overnight at 37⁰C. The next day, two additional aliquots of trypsin were added and incubated at 37⁰C for 4h each. At this stage, the protease digestion was evaluated on an aliquot of approximately 20 µg protein by SDSPAGE using a 10% NuPAGE polyacrylamide gel. After the digestion the samples were acidified with trifluoroacetic acid (TFA) (Thermo Fisher Scientific) to final concentration of 1% (v/v) and all insoluble material was removed by centrifugation.

#### Peptide purification

Two SepPak classic cartridges (Waters, Wat051910) were washed with 10 ml acetonitrile and then equilibrated with 6 ml 40% acetonitrile containing 0.1% TFA. Then the columns were washed and equilibrated with 0.1% TFA in water. The digestion products were passed through the columns to bind peptides, and the unbound material was reapplied. Each column was washed sequentially with 1-, 3- and 6 ml 0.1% TFA in water, and then the bound peptides were eluted three times with 2 ml 60% acetonitrile/0.1% TFA and the eluate was snap frozen and lyophilized.

To purify diGly-remnant peptides, we used the PTMScan Ubiquitin Remnant Motif Kit (CellSignaling catalogue number 5562) following the manufacturer’s instructions. Briefly, lyophilized peptides were dissolved in 1.4 ml immunoaffinity purification (IAP) buffer and the pH was adjusted to between 6 and 7 with 1 M Tris base, and then mixed by rotation for 2 h at 4°C with immunoprecipitation (IP) beads that had been washed previously 4 times in PBS. The IP beads were subsequently washed with IAP buffer and water, and bound peptides were eluted with 105 µl of 0.15% TFA in water and then lyophilized.

#### Mass spectrometry and processing of raw data

For MS analysis, peptides were dissolved in 0.1% TFA and loaded on a 50-cm Easy Spray PepMap column (75 μm inner diameter, 2 μm particle size, Thermo Fisher Scientific) equipped with an integrated electrospray emitter. Reverse phase chromatography was performed using the RSLC nano U3000 (Thermo Fisher Scientific) with a binary buffer system at a flow rate of 250 nl/min. Solvent A was 0.1% formic acid, 5% DMSO, and solvent B was 80% acetonitrile, 0.1% formic acid, 5% DMSO. The diGly enriched samples were run on a linear gradient of solvent B (2 - 40%) in 90 min, total run time of 120 min including column conditioning.

The Q Exactive was operated in data-dependent mode acquiring HCD MS/MS scans (R = 17,500) after an MS1 scan (R = 70, 000) on the 10 most abundant ions using MS1 target of 1 × 10^6^ ions, and MS2 target of 5 × 10^4^ ions. The maximum ion injection time utilized for MS2 scans was 120 ms, the HCD normalized collision energy was set at 28, the dynamic exclusion was set at 10 s, and the peptide match and isotope exclusion functions were enabled.

Protein and peptides were identified by the Andromeda search engine [71] integrated in the MaxQuant (Version 1.5.3.8) proteomics analysis software [26]. The protein database selected for the MS/MS searches contained *P. falciparum* protein sequences from PlasmoDB version 28 (http://plasmodb.org/common/downloads/release-28/) and human proteins from Uniprot (downloaded May 2016 https://www.uniprot.org/proteomes/UP000005640) supplemented with frequently observed contaminants. Andromeda search parameters for protein identification were set to a precursor mass tolerance of 20 ppm for the first search and 6 ppm for the main search. The mass tolerance for MSMS fragmentation spectra was set at 0.5 Da. Trypsin protease specificity was enabled allowing up to 4 mis-cleaved sites. Di-glycine modification of lysine, deamination of glutamine and asparagine, oxidation of methionine, and protein N-terminal acetylation were set as variable modifications. Carboxyamidomethylation of cysteine was specified as a fixed modification. The minimal required peptide length was specified at 6 amino acids. MaxQuant was used to perform internal mass calibration of measured ions and peptide validation by the target decoy approach. Peptides and proteins with a false discovery rate (FDR) lower than 1% were accepted. Only ubiquitination sites with a localization probability >0.75 in one of the life cycle stages were accepted.

### Bioinformatic analysis

#### Ubiquitylation motif analysis

Ubiquitylation motifs were identified using MotifX [72] with the following parameters: motif window = 13 amino acids, *P. falciparum* PlasmoDB version 28 proteins (http://plasmodb.org/common/downloads/release-28 /) as background, statistical significance threshold : p-value < 1 * 10^-6^, fold motif increase ≥ 2, and a motif frequency > 5 Sequence logos were generated with Weblogo 3 (http://weblogo.threeplusone.com/create.cgi).

#### Gene ontology (GO) analysis

Gene ontology (GO) analyses were performed with the software package Ontologizer [73], with the gene association components downloaded from http://www.geneontology.org: gene ontology. Overrepresented GO terms for the merozoite ubiquitome and intracellular ubiquitome relative to the background of the *P. falciparum* proteome (∼5500 proteins) were computed by the parent−child union approach and corrected for multiple testing by the Benjamini and Hochberg method and were considered significant for adjusted p-values lower than 0.05. The results were visualised in an enrichment map with the Cytoscape plugin Enrichment Map v1.2 [74].

#### Pathway enrichment analysis

Enrichment for Malaria Parasite Metabolic Pathways (MPMP) [28] (http://mpmp.huji.ac.il/) pathway terms of the merozoite unbiquitome relative to all 5500 *P. falciparum* proteins was determined by a one-tailed Fisher’s exact test (FET) adjusted for multiple testing correction by the Benjamini-Hochberg method.

### PfUBA1 homology model, purification of recombinant protein and analysis of activity

#### UBA1 homology model

Using the published structure of ScUBA1 in complex with MLN7243 covalently linked to the C-terminus of ubiquitin (PDB: 5LDJ) a homology model of the PfUBA1-ubiquitin-MLN7243 complex was generated using MOE [Molecular Operating Environment (MOE), 2018.01; Chemical Computing Group ULC, 1010 Sherbrooke St. West, Suite #910, Montreal, QC, Canada, H3A 2R7, 2018]. The structure was corrected and protonated using the Protonate3D tool. The sequences of PfUBA1 and ScUBA1 were aligned, resulting in an overall identity of 39.6% and an identity of 78.6% for the binding pocket, defined as residues lying within 5 Å of the inhibitor. The homology model was prepared using the induced fit option with MLN7243 used as environment. The model was checked for correct protein geometry and incorrect loops were remodelled. The final model was energy minimized using the force fields AMBER10FF for the protein and MMFF94x for MLN7243. The homology model was validated with PROCHECK [75] resulting in a structure in which 73.0% of the residues are located in the most favoured regions, 24.4% in additionally allowed regions, 1.4% in generously allowed regions and 1.2% of the residues in disallowed regions of the Ramachandran diagram. Residues lying in the disallowed regions do not belong to the binding site and are located in outer loop regions of the model.

#### Expression and purification of recombinant UBA1

A synthetic codon-optimised *uba1* gene (Supplementary Figure 4) corresponding to amino acid residues Thr39 to Lys1140 was designed and purchased from GeneArt, amplified by PCR and inserted into the pIEX/Bac-3 insect cell expression vector (Novagen) using ligation independent cloning. The insert sequence was verified by DNA sequencing. This vector produces a protein with an N-terminal HRV 3C cleavable His_10_-tag fusion (MA**HHHHHHHHHH**GA<u>LEVLFQGP</u>G, where the sequence in bold is the His_10_-tag and the underlined sequence is the HRV 3C site).

#### Protein Expression

For protein expression in insect cells, recombinant baculovirus was prepared by co-transfecting *sf9* cells, cultured at 28°C in SF900 III serum-free medium (Invitrogen), with pIEX/Bac3-*pf*UBA1 plasmid and linearised flashbac bacmid (Oxford Expression Technologies) according to the manufacturer’s instructions. Expression of the *Pf*UBA1 protein on a small scale was tested by infecting a 25 ml culture of insect cells (∼2 x 10^6^ cells/ml) with 100 µl of high titre virus stock and cultivation for 48 h. The cells were pelleted at 3,000 rpm at 4°C, and suspended in 7 ml of 50 mM Tris-HCl pH 7.5, 500 mM NaCl, 10 mM MgCl_2_, 10% glycerol, 10 mM imidazole pH 8.0, 1mM TCEP, 5U universal nuclease (Pierce) and protease inhibitors (Roche complete, EDTA Free). Lysed cells were clarified at 5,000 rpm at 4°C. The supernatant was loaded onto a 100 µl Talon resin column (Clontech-Laboratories) and protein purification was carried out following the manufacturer’s instructions.

#### Protein purification

Large scale expression was carried out in roller bottles by infecting 600ml culture containing ∼2 x 10^6^ cells/ml per roller bottle with 2.4 ml of high titre virus. Cells were harvested 48h post infection and resuspended in 25 mM NaHCO_3_. An equal volume of 2x lysis buffer containing 100 mM Tris-HCl pH 7.5, 1 M NaCl, 20 mM imidazole pH 8.0, 10% glycerol, 20 mM MgSO_4_, 2 mM TCEP, 1x protease inhibitor cocktail and 1:1000 universal nuclease was added before sonication. The lysate was centrifuged at 55,000g for 30 min at 4 °C. The supernatant containing solubilized His_10_-pfUBA1 was loaded on 5 ml of Talon resin and washed with wash buffer containing 50 mM Tris-HCl pH 7.5, 300 mM NaCl, 5% glycerol, 1 mM TCEP, and 15 mM imidazole. Bound proteins were eluted in wash buffer containing 250 mM imidazole. Pooled fractions were diluted 1:1 with 50 mM Tris-HCl pH7.5 and loaded onto a 1 ml MonoQ column (GE Healthcare) pre-equilibrated in buffer A (50 mM Tris -HCl pH 7.5, 150 mM NaCl, 0.5 mM TCEP) at 4°C. Bound *Pf*UBA1 was eluted with a linear gradient to 100% buffer B (50 mM Tris -HCl pH 7.5, 1 M NaCl, 0.5 mM TCEP). Fractions containing PfUBA1 were pooled, concentrated to 100 µM, flash-frozen in liquid nitrogen and stored at -80°C.

#### Thermal Stability Measurements

The thermal unfolding profile of purified *pf*UBA1 was measured using nanoDSF. Standard, nanoDSF grade Prometheus NT.48 capillaries (NanoTemper Technologies) were filled with ∼10 μL measurement sample at 100 µM. Back reflection (light scattering) and nanoDSF (nano-differential scanning fluorimetry) measurements were performed using a Prometheus NT.48 (NanoTemper Technologies) at 5% excitation power and with a thermal ramp from 25 to 95°C with a heating rate of 2.5 °C/min. Fluorescence data for protein unfolding were analysed using the GraphPad Prism software. The fluorescence (F) ratio F330nm/F350nm was used for inflection point determination, which represents the melting temperature (Tm). Unfolding transition temperatures (T_m_) were automatically determined by the software using the Boltzman equation.

#### In vitro analysis of the activation, transthioesterification and inhibition of UBA1

A ubiquitin cysteine mutant (M1C) was labelled with Cy5 maleimide mono-reactive dye (GE Healthcare) as described [76]. E2s were expressed and purified as described [76, 77]. In activation and transthioesterification assays 30 nM Cy5-Ub was incubated with 1 µM E1 (*Pf* UBA1 or *Mm* Ube1) in reaction buffer (50 mM HEPES pH 7.5, 150 mM NaCl, 5 mM MgCl_2_) at room temperature, and the Ub activation and thioester formation was started by adding 100 µM ATP. After 5 min 10 µM E2 (either UbcH5, UbcH7, Cdc34 or Ubc13) was added to aliquots of the reaction mixture and incubated for further 5 min. Two samples were taken at every step and immediately mixed with 2 x LDS sample buffer (NuPAGE, Thermo Fisher) minus and plus 500 mM DTT. Samples were analysed by SDS-PAGE on 4-12% gradient gels (NuPAGE, Thermo Fisher). Gels were scanned using a Typhoon FLA 9500 scanner (GE Healthcare) to visualize the Cy5-Ub containing bands.

In E1 inhibition assays 30 nM Cy5-Ub and 1 µM UBA1 were incubated with a two-fold serial dilution of the inhibitor (MLN7243 or MLN4924), from 40 µM to 0.039 µM, in reaction buffer with 2% DMSO. 100 µM ATP was added and the reaction mixtures were incubated for 30 min at room temperature before they were mixed with 2 x LDS sample buffer without DTT. Samples were analysed by SDS-PAGE and gels scanned as described above. Band intensities were quantitated using the ImageQuant software (GE Healthcare). The percentage intensities of the Cy5-Ub-E1 thioester band was plotted versus inhibitor concentration. IC_50_ values were determined by non-linear least square fitting using the Hill equation in Prism 7 (GraphPad).

## Acknowledgements

This work was supported. by the Francis Crick Institute which receives its core funding from Cancer Research UK (FC001097), the UK Medical Research Council (FC001097), and the Wellcome Trust (FC001097). AP was funded by a British Council Newton Fund PhD Placement. SB and NT were supported by fellowships from GK 2243 funded by the Deutsche Forschungsgemeinschaft. BS was supported by a Biotechnology and Biological Sciences Research Council Project grant (BB/R003750/1). We gratefully acknowledge Dr Hagai Ginsburg for providing MPMP data enabling the pathway enrichment analysis.

## Database deposition

The mass spectrometry proteomics data have been deposited to the ProteomeXchange Consortium via the PRIDE [78] partner repository with the dataset identifier PXD014998.

