## Supplementary Figures 1 to 5 and Table 8 for "Protein ubiquitylation is essential for the schizont to merozoite transition in *Plasmodium falciparum* blood-stage development"

**Supplementary information**

Yang Wu[1], Vesela Encheva[2], Judith L. Green[1], Edwin Lasonder[3], Adchara Prommaban[1,4], Simone Kunzelmann[5], Evangelos Christodoulou[5], Munira Grainger[1], Ngoc Truongvan[6], Sebastian Bothe[7], Vikram Sharma[4], Wei Song[8], Irene Pinzuti[8], Chairat Uthaipibull[9], Somdet Srichairatanakool[4], Veronique Barault[10], Gordon Langsley[11], Hermann Schindelin[6], Benjamin Stieglitz[8], Ambrosius P. Snijders[2], Anthony A. Holder[1*]

[1]Malaria Parasitology Laboratory, [2]Mass Spectrometry Proteomics, [5]Structural Biology Science Technology Platform, [10]Translation, The Francis Crick Institute, London NW1 1AT, UK; [3]School of Biomedical Science, Faculty of Medicine & Dentistry University of Plymouth, Plymouth PL6 8BU, UK; [4]Department of Biochemistry, Faculty of Medicine, Chiang Mai University, Chiang Mai 50200, Thailand; [6]Rudolf Virchow Center for Experimental Biomedicine, Universität Würzburg, 97070 Würzburg, Germany; [7]Institute of Pharmacy and Food Chemistry, Department of Chemistry and Pharmacy, University of Würzburg, 97074 Würzburg, Germany; [8]School of Biological and Chemical Sciences, Queen Mary University of London, London E1 4NS, UK; [9]National Center for Genetic Engineering and Biotechnology (BIOTEC), Khlong Nueng, Khlong Luang, Thailand; [11]Laboratoire de Biologie Cellulaire Comparative des Apicomplexes, Institut Cochin, Faculté de Medecine, Université Paris Descartes, 75014 Paris, France.

**Supplementary figures**

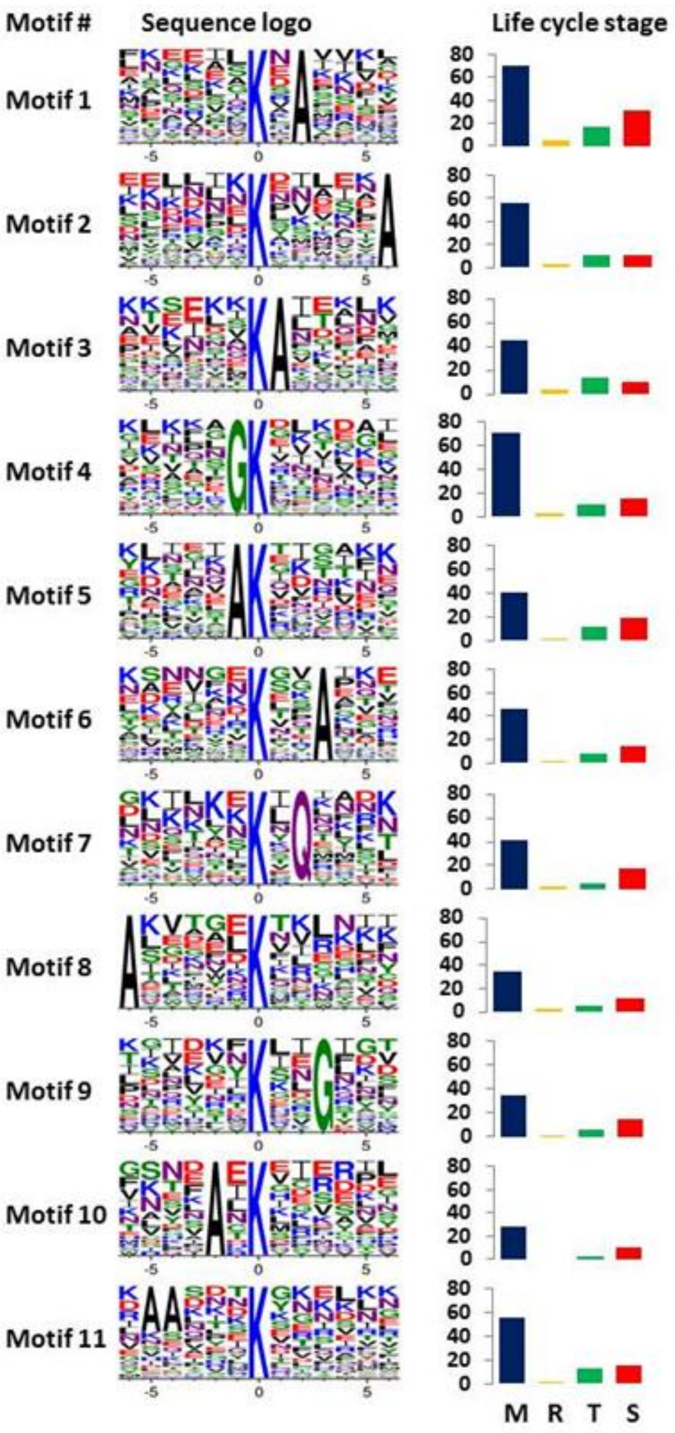


Supplementary Figure S1: Sequence logo depiction of the 11 most abundant ubiquitylated motifs (covering 644 out of 1464 sites) detected by Motif X analysis.

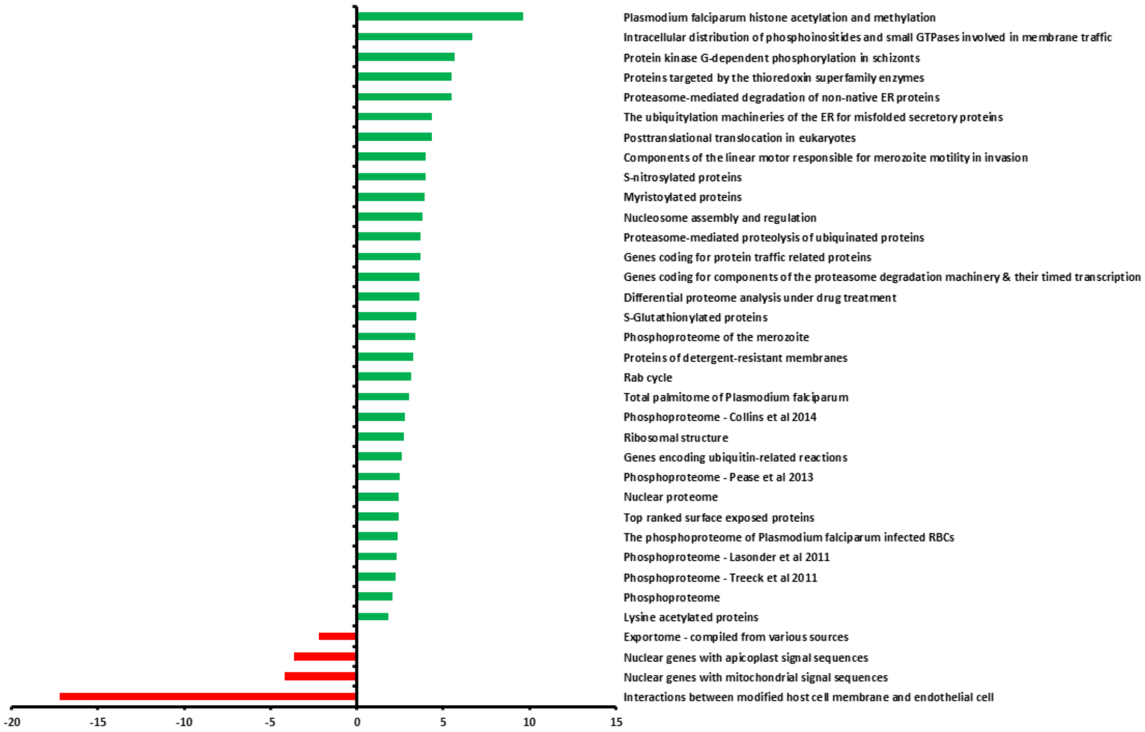


Supplementary Figure S2. MPMP pathway enrichment analysis of the merozoite ubiquitome. Significantly upregulated pathways are in green and downregulated pathways in red.

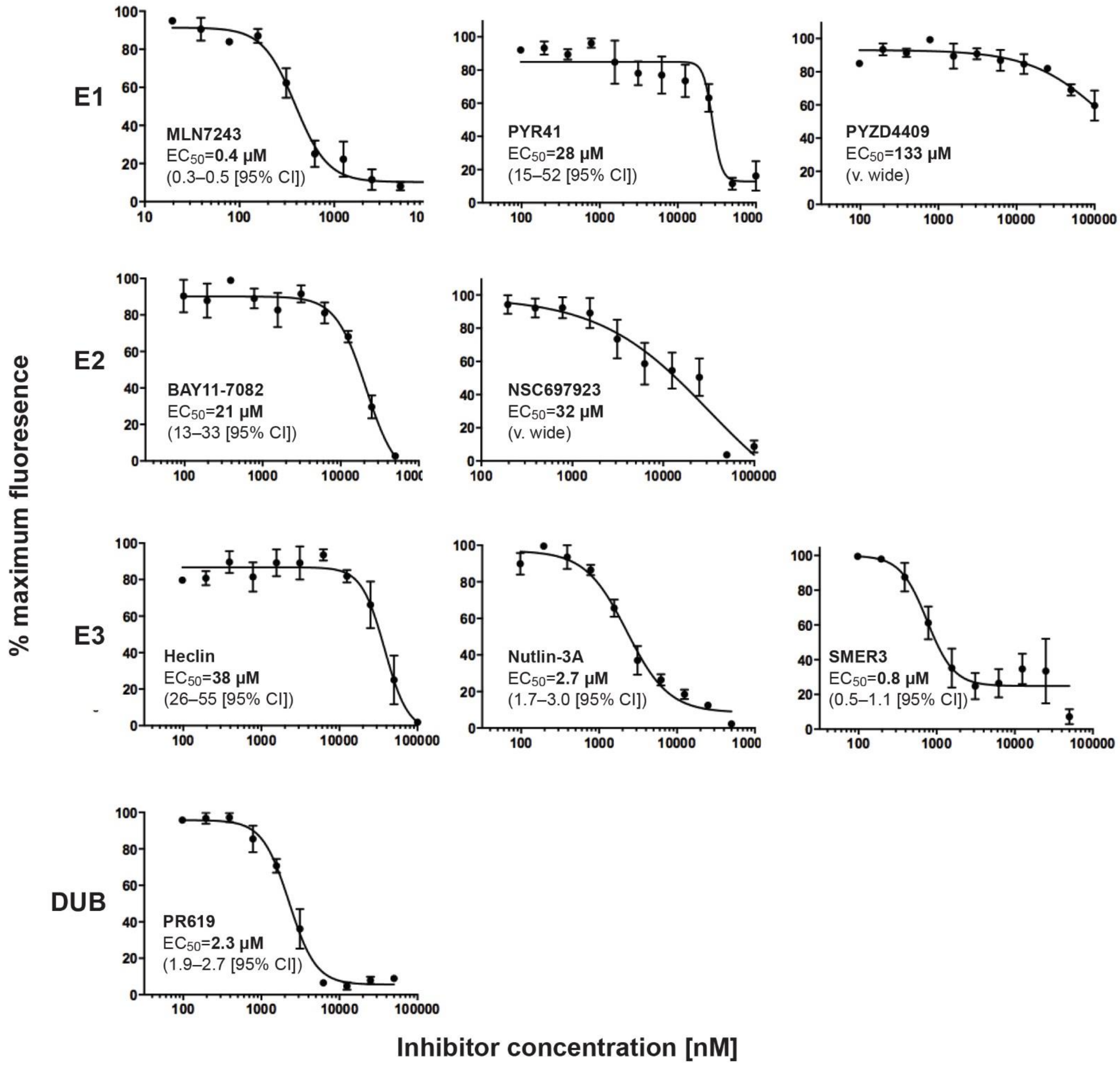


Supplementary Figure S3: Ability of commercially available inhibitors of the ubiquitylation pathway to inhibit parasite growth in vitro. For each inhibitor an $EC_{50}$ is calculated from the curve.

```
        T E N K I D T D L Y S R Q L G T Y G F D L M N K L V K L N I L I I N V K G V G L E C A K N L I L S G P
1       GGATCCAACCGAGAACAAAATTGATACCGATCTGTATAGCCGTCAGCTGGGCACCTATGGTTTTGATCTGATGAATAAACTGGTGAAACTGAACATCCTGATCATCAATGTTAAAGGTGTTGGTCTGGAATGCGCCAAAAATCTGATTCTGAGCGGTCCG

        Q S V C I Y D N D I C D I S D I G V N F Y I N E K D V E D K S C R S D A V L K E L Q E L N N Y V H I Y N Y
161     CAGAGCGTTTGTATTTATGATAATGATATTTGCGACATCAGCGACATCGGCGTGAACTTTTATATCAACGAAAAAGACGTGGAAGATAAGAGCTGTCGTAGTGATGCAGTTCTGAAAGAACTGCAAGAACTGAACAACTACGTGCACATCTATAACTATA

        K G T I E K N W L E N F D V V I C C D I N K E D L I K Y N N M I R G I D K K R I A F L S C N I Y G L C G Y I

321     AAGGCACCATCGAAAAGAACTGGCTGGAAAATTTTGATGTGGTGATTTGCTGCGATATCAACAAAGAGGATCTGATCAAGTATAACAACATGATCCGTGGCATCGACAAAAAACGTATTGCATTTCTGAGCTGCAACATTTATGGTCTGTGCGGTTATAT

        F V D F N K E F I C Y D S N G E Q V K S C N V S K I S K E L E G K V S F D F D K T S P F E E G D Y V Q F S
481     CTTCGTGGACTTCAATAAAGAGTTCATCTGCTACGATAGCAATGGCGAACAGGTTAAAAGCTGCAATGTTAGCAAAATCAGCAAAGAGCTGGAAGGCAAAGTCAGCTTTGATTTTGATAAAACGAGCCCGTTTGAAGAGGGTGATTATGTGCAGTTTAGC

        N V E G M T E I N N K I Y K I K N L K K Y T F E I G D T S L Y S E Y I K G G I C T Q V K K H L K L N F Y P
641     AATGTTGAAGGCATGACCGAAATCAACAACAAAATCTATAAAATCAAAAACCTGAAAAAGTACACCTTCGAGATTGGTGATACCAGCCTGTATAGCGAATATATCAAAGGTGGTATTTGCACCCAGGTGAAAAAACATCTGAAGCTGAATTTCTATCCGT

        Y E Y I C V N P L N N E N I S N N E Q K H N Q N D N H F L D T C N N I I Y E N I P Q P N S F I I S D Y A K F
801     ACGAGTATATTTGCGTGAATCCGCTGAATAACGAAAACATCAGCAACAACGAACAGAAGCATAACCAGAACGATAACCATTTTCTGGATACCTGCAACAACATCATCTATGAAAACATTCCGCAGCCGAACAGCTTTATCATTAGCGATTATGCCAAATT

        D M S N H L H Y S I Q A L K W Y E L Q N E K G L P E N S D E D A L E K I Y N Y A V T L N N K D K E E K K S
961     TGATATGAGCAACCATCTGCACTATAGCATTCAGGCACTGAAATGGTATGAACTGCAGAATGAAAAAGGTCTGCCGGAAAATAGTGATGAAGATGCCCTGGAAAAAATCTACAATTATGCAGTGACCCTGAACAACAAGGACAAAGAAGAGAAAAAAAGC

        Y A V E Q L K K D V V Y N V C R Y S K S H I A P V A S F F G G L L A Q E V I K F T G K Y M P I Y Q L L Y L
1121    TACGCAGTGGAACAGCTGAAAAAAGATGTTGTGTATAATGTGTGCCGCTACAGCAAAAGCCATATTGCACCGGTTGCAAGCTTTTTTGGTGGTCTGCTGGCACAAGAAGTGATCAAATTTACCGGCAAATATATGCCGATTTACCAGCTGCTGTATCTGG

        D F F E C I S L N E K V D I N E I K K M N C K N D N I I T V F G K S F Q K K L N N L N V F L V G S G A L G C
1281    ATTTTTTTGAATGCATTAGCCTGAACGAGAAGGTGGATATTAACGAGATCAAAAAGATGAATTGCAAAAACGATAACATCATCACCGTGTTCGGTAAAAGCTTCCAGAAAAAACTGAATAACCTGAACGTGTTTCTGGTTGGTAGCGGTGCACTGGGTTG

        E Y A K L F S L L D M C T R N S E Q N T N L N Q N N I D N N L A C C G K L T I T D N D N I E V S N L N R Q
1441    TGAATATGCAAAACTGTTTAGCCTGCTGGATATGTGTACCCGTAATAGCGAACAGAATACCAATCTGAACCAGAACAACATCGATAACAATCTGGCATGTTGTGGCAAACTGACCATTACCGATAATGACAATATTGAGGTGAGCAATCTGAATCGCCAG

        F L F R R E H V G K S K S L V S S E I I K K K N N N M H V Q S L E T K V G A E N E H I F N E E F W T K Q N
1601    TTTCTGTTTCGTCGTGAACATGTGGGTAAAAGCAAAAGTCTGGTTAGCAGCGAGATCATCAAAAAAAAGAACAACAACATGCACGTCCAGAGCCTGGAAACCAAAGTTGGTGCAGAAAATGAACACATCTTCAACGAAGAATTCTGGACCAAACAGAACA

        I I V N A L D N I Q A R Q Y V D N K C V W Y S K P L F E S G T L G T K G N V Q V I I P Y L T Q S Y N D S Y D
1761    TTATTGTGAACGCCCTGGATAATATTCAAGCCCGTCAGTATGTTGATAACAAATGCGTTTGGTATAGCAAACCGCTGTTTGAAAGCGGCACCCTGGGCACCAAAGGTAATGTTCAGGTGATTATTCCGTATCTGACCCAGAGCTATAATGATAGCTATGA

        P P E D S I P L C T L K H F P Y D I V H T I E Y A R D I F Q G L F Y N T P L S I K Q F L N D K E E Y I N K
1921    CCCTCCGGAAGATAGCATTCCGCTGTGTACCCTGAAACATTTTCCGTATGATATTGTGCACACCATTGAATATGCCCGTGATATTTTTCAGGGCCTGTTTTATAACACACCGCTGAGCATCAAACAGTTTCTGAACGATAAAGAAGAGTACATCAACAAG

        I Q E E G N N A S L L E N L Q N V I N S L K E I S S Q C N F D F C I K K S V E L F H N N F I N Q I N Q L L
2081    ATTCAAGAAGAAGGTAATAACGCCAGTCTGCTGGAAAACCTGCAGAACGTGATTAATAGCCTGAAAGAAATTAGCAGCCAGTGCAACTTTGACTTCTGCATTAAAAAGTCCGTGGAACTGTTTCACAACAACTTCATCAATCAGATTAATCAACTGCTGT

        Y S F P L D Y K L S S G E Y F W V G Q K K P P Q P I V F D V N N E M I Q E F L L S T S N L L A Q V Y N I P P
2241    ATAGCTTCCCGCTGGATTATAAACTGAGCAGCGGTGAATATTTCTGGGTGGGCCAAAAAAAACCGCCTCAGCCGATTGTTTTTGACGTGAACAATGAAATGATCCAAGAATTTCTGCTGAGCACCAGCAATCTGCTGGCCCAGGTTTATAACATTCCTCC

        C F D I N Y I I N V A K K I E V K P F E P K K V K I N M D E K N L N N I S I S F A E E E K I I D D F C K E
2401    GTGTTTCGATATTAACTATATCATCAACGTGGCGAAAAAAATCGAGGTGAAACCGTTTGAACCGAAAAAGGTGAAAATCAACATGGACGAGAAGAACCTGAATAACATTAGCATTAGCTTTGCCGAGGAAGAGAAGATCATTGACGATTTTTGTAAAGAG

        L L N I P T N N I K I N P I E F D K D E Q T N L H V N F I Y A F S N L R A I N Y K I N T C D K L K A K I V
2561    CTGCTGAATATCCCGACCAACAACATTAAAATCAACCCGATCGAGTTCGATAAAGATGAACAGACCAATCTGCATGTGAACTTCATTTATGCCTTTAGTAATCTGCGTGCCATCAACTATAAGATCAATACCTGCGATAAACTGAAGGCCAAAATTGTTG

        A G K I I P A L A T T T S I I T G L V G I E L L K Y V N Y Y D N I Q A Y V K L S D E Q R K K E K H D V L S Y
2721    CGGGTAAAATCATTCCGGCACTGGCAACCACCACCAGTATTATTACCGGTCTGGTGGGTATTGAACTGCTGAAATATGTGAACTACTACGACAACATTCAGGCCTATGTTAAACTGAGTGATGAGCAGCGCAAAAAAGAAAAGCATGATGTTCTGAGCTA

        F K N A F I N S A L P L F L F S E P M P P L R M M D K E Y D E L M K G P V K A I P N G F S S W D K I V I S
2881    CTTCAAGAACGCCTTTATTAACAGCGCACTGCCGCTGTTTCTGTTCAGCGAACCGATGCCTCCGCTGCGTATGATGGATAAAGAATATGATGAACTGATGAAAGGTCCGGTTAAAGCAATTCCGAATGGTTTTAGCAGCTGGGATAAAATTGTGATCAGC

        I K N G T I K D L I D H I N E K Y S I D V N L I S V G N A C L Y N C Y L P A H N K E R L N K P I H E L Y K
3041    ATCAAAAACGGCACCATCAAAGACCTGATTGATCACATCAATGAGAAGTATAGCATCGACGTGAATCTGATTAGCGTTGGTAATGCCTGTCTGTATAATTGTTATCTGCCTGCGCATAACAAAGAGCGTCTGAATAAACCGATTCACGAACTGTATAAGC

        Q I S K Q D L L E D K N Y I I V E A S C S D Q D L V D V L I P S I Q F I Y K *
3201    AGATCAGTAAACAGGACCTGCTTGAGGATAAAAACTACATTATCGTTGAAGCCAGCTGTAGCGATCAGGATCTGGTGGATGTGCTGATTCCGAGCATCCAGTTTATCTACAAATGATAGTAACTCGAG
```

Supplementary Figure S4: Synthetic gene for the expression of PfUBA1 (PF3D7_1225800), coding for amino acid residues 39 to 1140 (the C-terminus of the protein)

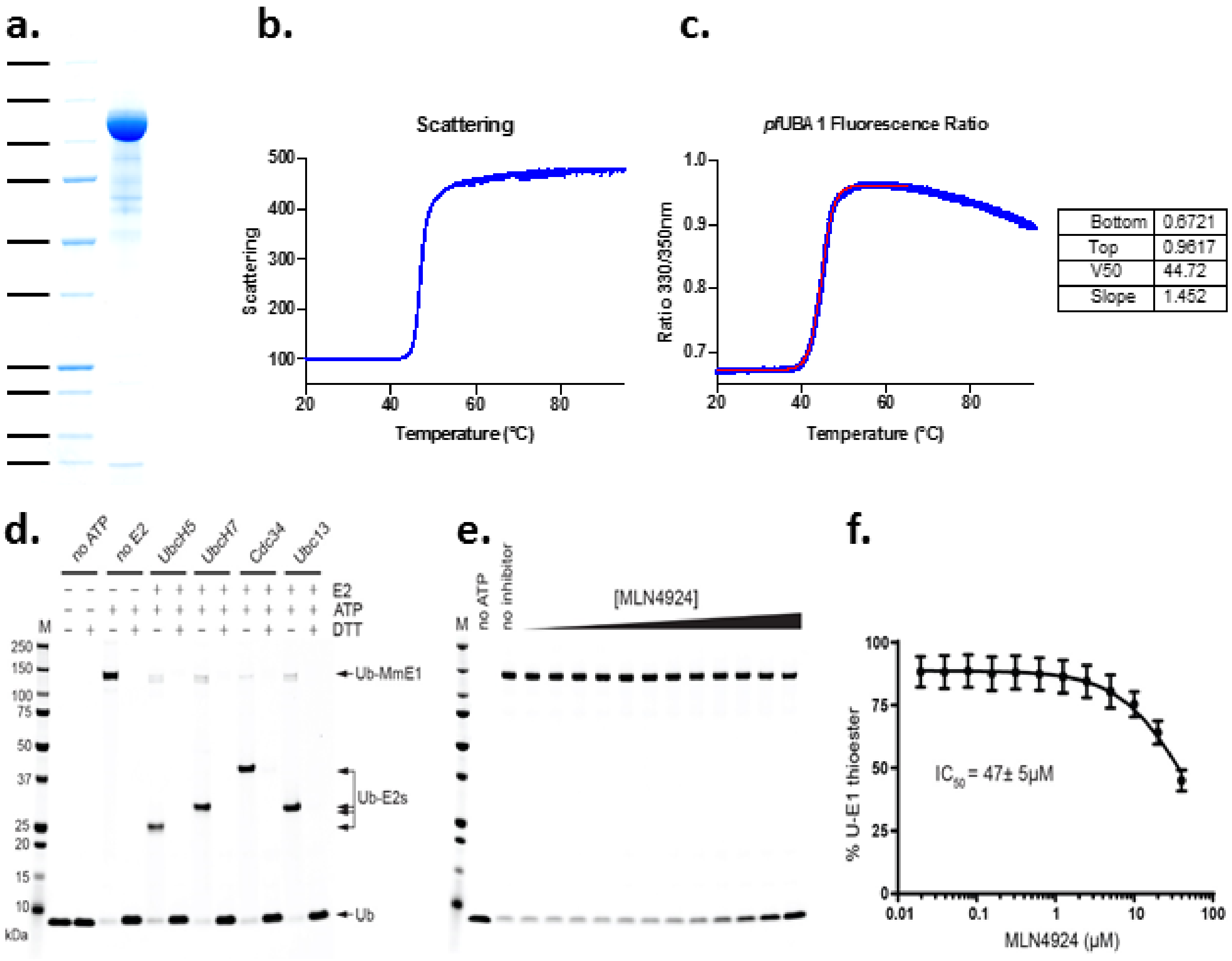


Supplementary Figure S5. Recombinant enzyme and properties. (a) Purification and stability of recombinant PfUBA1. The protein was expressed in the baculovirus-insect cell system and purified on Talon and MonoQ resins; analysis of final product by Coomassie blue stained SDS-PAGE. The thermal stability of the protein was analysed by (b) light scattering and (c) fluorescence to identify the melting temperature. (d) Validation of the in vitro thioesterification and transthioesterification assays: *Mus musculus* UBA1 is thioesterified by fluorescently labelled ubiquitin in the presence of ATP, and transthioesterifies various human E2s. (e) the lack of activity of a Nedd8 E1 inhibitor against PfUBA1; MLN4924, added in doubling dilutions, is a poor inhibitor of PfUBA1 thioesterification. (f) The data from panel (e) enabled the $IC_{50}$ of MLN4924 to be calculated.

**Supplementary Tables**

| | |
|---|---|
| Supplementary Table 1 | All the parasite peptides and proteins ubiquitylation data (Excel file) |
| Supplementary Table 2 | All the human peptides and proteins ubiquitylation data (Excel file) |
| Supplementary Table 3 | Numbers of sites per protein distribution (Excel file) |
| Supplementary Table 4 | Listing of GO terms (Excel file) |
| Supplementary Table 5 | Pellicle proteins (Excel file) |
| Supplementary Table 6 | exported proteins (Excel file) |
| Supplementary Table 7 | Histones (Excel file) |

Supplementary Table 8. Oligonucleotides used in this study

| PRIMER | SEQUENCE | |
|---|---|---|
| 107for | ATTGTGATCGATTAAATCTTTAA | Guide 107 |
| 107rev | AAACTTAAAGATTTAATCGATCA | |
| 109for | ATTGTTATTTGCCTGCACACAACA | Guide 109 |
| 109rev | AAACTGTTGTGTGCAGGCAAATAA | |
| 111for | ATTGTTAAGACATCTACGAGGTCT | Guide 111 |
| 111rev | AAACAGACCTCGTAGATGTCTTAA | |
| 113 | gatcCCGCGGTGATGAGTGCACAAATATATATATATATATAT | |
| 114 | gatcAAGCTTCAAAACGGTGATGGGTATATATACTTCCTTGTT | |
| 116 | GTTCATATAAACATCAAAATGAATCAACAATGATTC | screening primers |
| 121 | ATATTTCCATATCCTTTGCCGAAGAAGAAAAAA | |
| GFPrev | CTCCAGTGAAAAGTTCTTCTCC | |
| HArev | CCTTTACCGCGGTCAAGCGTAAT | |
